# Identification of a meiosis-specific chromosome movement pattern induced by persistent DNA damage

**DOI:** 10.1101/2020.07.23.218016

**Authors:** Daniel León-Periñán, Alfonso Fernández-Álvarez

## Abstract

As one of the main events occurring during meiotic prophase, the dynamics of meiotic chromosome movement is not yet well understood. Currently, although it is well-established that chromosome movement takes an important role during meiotic recombination promoting the pairing between homologous chromosomes and avoiding excessive chromosome associations, it is mostly unclear whether those movements follow a particular fixed pattern, or are stochastically distributed. Using *Schizosaccharomyces pombe* as a model organism, which exhibits dramatic meiotic nuclear oscillations, we have developed a computationally automatized statistical analysis of three-dimensional time-lapse fluorescence information in order to characterize nuclear trajectories and morphological patterns during meiotic prophase. This approach allowed us to identify a patterned oscillatory microvariation during the meiotic nuclear motion. Additionally, we showed evidence suggesting that this unexpected oscillatory motif might be due to the detection of persistent DNA damage during the nuclear movement, supporting how the nucleus also regulates its oscillations. Our computationally automatized tool will be useful for the identification of new patterns of nuclear oscillations during gametogenesis.

## INTRODUCTION

Meiosis is an essential process regarding the improvement of genetic diversity, allowing the generation of new allelic combinations. It consists of two consecutive rounds of nuclear divisions, known as meiosis I (MI) and meiosis II (MII), after a single round of DNA replication, which ensure the correct distribution of chromosomes to haploid gametes from diploid parental cells ^1^. During meiotic prophase, chromosome oscillations driven by cytoskeleton forces aim to move the chromosomes in order to promote the recognition and pairing between homologous ^2–4^. Recently, it has also been observed that meiotic nuclear movements allow the elimination of unwanted chromatin to nuclear envelope (NE) associations and excessive inter-chromosomal entanglements ^5^. Despite the ubiquity of meiotic nuclear movement throughout evolution, the key question about if these oscillations follow a fixed pattern of movement or are stochastic has not been disclosed yet. To address this point, we have used the fission yeast *Schizosaccharomyces pombe* in which during prophase, the nucleus displays vigorous and continuous movements across poles lasting nearly 2 hours before MI. Due to the morphology acquired by the nucleus, this stage is known as *horsetail* movements (Supplementary Figure S1A). In particular, it has also been demonstrated that homologous chromosome pairing-unpairing dynamics is depending on the nuclear extension towards poles (*horsetail* shape) or the centre (rounded shape) ^6^. Hence, in fission yeast as well as in most of the eukaryotes, nuclear oscillations are essential for faithful chromosome recombination and segregation during meiosis ^7–9^.

In *S. pombe*, meiotic nuclear movements are preceded by the telomere bouquet formation, a meiotic prophase-specific chromosomal configuration where the telomeres cluster together in a region of the NE, frequently, near the centrosome or Spindle Pole Body (SPB, centrosome equivalent in yeast) ^10–12^. This telomere-NE interaction is maintained by Bqt1 and Bqt2 proteins, two meiotic-specific proteins that form a bridge between the telomeric proteins Rap1 and Taz1, and the SUN-domain protein Sad1 ^13,14^. Meiotic prophase is characterized by prominent oscillations of the SPB driven by microtubules (MTs) and, thanks to the telomere bouquet formation, chromosomes can follow the SPB movement adopting the mentioned above *horsetail* shape ^2,15^. In more detail, SPB oscillations are driven by the polymerization dynamics of cytoplasmic astral MTs, mainly constituted by proteins such as alpha tubulin-2 (Atb2), and the self-organization of motor proteins such as dynein, composed by both Dlc1 and Dhc1(light and heavy chains), with the contribution of diffusive and directed motions ^16^. To exert forces that drive nuclear migration, dynactin, the dynein activator complex, is required to be anchored to the cell cortex through cortical factor Num1 when pulling MTs, as well as to other dynactin subunits ^17,18^. On the other hand, organization of astral MT arrays takes place thanks to the role of Hrs1, a meiosis-specific coiled-coil protein which localizes to the outer side of the SPB, directly interacting with Alp4, a component of the gamma-tubulin ring complex (γ-TuRC) ^19,20^ (Supplementary Figure S1B).

There are two main mechanistic models in the literature that might explain how the oscillatory behaviour is maintained: i) the SPB stops after supplying biochemical signals when reaching cell poles, deactivating the anchoring of dynein at a particular pole ^7^. Then, SPB oscillations are cyclically re-established when MTs reach the cortex at the opposite pole. ii) On the other hand, Vogel *et al*. proposed the *minimal model* ^21^. As experimental evidence clarifies, such as from single-molecule imaging ^22^; the *minimal model* suggests that the own MT polymerization and dynein diffusion dynamics are the two central regulators of the oscillatory patterns:. as fewer dyneins remain on the trailing side, they begin to feel higher loads that lead to its detachment from the microtubule: this establishes an asymmetric distribution across it, promoting the oscillatory mechanism. Besides, while not bound to cargo, dyneins remain in a non-motile state ^23^. It was consequently checked by laser ablation experiments that artificially directed MT breakdown changes the direction of the displacement, as predicted by the model ^21^. Briefly, oscillations are the result of a pulling force exerted by anchored dyneins, with a lower number of anchored dyneins on the trailing side, being lateral MT-dynein anchoring enough. However, there are still critical open questions like the cause of the slowdown or pausing of the SPB displacement when reaching cell tips ^24^. Finally, no model considers the full integration and stage transition during prophase concerning other cellular processes. For all these reasons, there is still a lack of knowledge about the molecular rules behind meiotic nuclear oscillations, a mechanism of enormous relevance for gametogenesis.

In this work, we have gained information about the behaviour of meiotic nuclear movement and present a model for the integration of horsetail oscillations within prophase, in terms of nuclear morphology and trajectory, also considering cellular responses that can induce alterations of the oscillatory mechanism. <u>Imaging</u> and software pipelines (Supplementary Figure S1C) have been developed and applied to assess this phenomenon as the sum of smaller parts or motifs that, to the best of our knowledge, have not been previously described. Using our approach, we identified new patterns during the nuclear oscillations and proposed that they might define a mechanism to resolve persistent DNA damage. We think that our tool will be useful for future studies focused on nuclear dynamics in yeast and other model organisms for gametogenesis studies.

## RESULTS

### 2D nuclear morphology and trajectory in meiotic prophase are three-segmented

Current models divide the meiotic prophase in fission yeast, defined by the formation and disassembly of the telomere bouquet, into two phases: i) *horsetail stage*, characterized by the most intense nuclear oscillations; and ii) *post-horsetail stage*, a short segment at meiosis onset where the telomere bouquet is disassembled without nuclear movement ^25^. To further characterize meiotic nuclear movement and identify possible patterns or random behaviours in the trajectory and morphology of the nucleus, we developed an imaging and software pipeline. With this aim, we applied our approach to a fission yeast *wt* strain harbouring the GFP-endogenously tagged Sid4 (an integral component of the SPB) and Hht1-CFP (endogenous tagging of the one copy of histone 3 variant) to visualize chromosome behaviour.

Three-dimensional blobs representing SPB were successfully obtained with our live-microscopy settings for each frame in all time-lapse experiments, as well as their XYZ coordinates (see Methods) (Figure 1A-B). Interestingly, LOWESS fitting and Lavielle segmentation with optimum parameters (Supplementary Figure S2) revealed that meiotic prophase was successfully segmented for the XY differenced case, obtaining three significant segments instead of the current model suggesting two. Hence, stages were named according to the dominant frequencies and amplitudes as *high nuclear motion, medium nuclear motion* and *low nuclear motion*, respectively (Figure 1C-D). Coefficients obtained for ARIMA analysis of the residuals for each segment of the three-dimensional displacements show that displacement along X and Y axis both have a periodicity near to 5 lags. Remarkably, the SPB movement for the ZX coordinates was also explored in our analysis, finding no significant segmentation. Displacement along all Z-axis has values near to random walk models, thus indicating for the first time that nuclear motion in Z-axis follows a random trajectory (Figure 1E and Supplementary Figure S3); for this reason, we discarded Z-axis data for further analysis. In more detail for each of the three newly identified stages, ARIMA analysis significantly revealed that: (i) *high nuclear motion* stage had a lower periodicity than *medium nuclear motion* phase and (ii) *low nuclear motion* segment behaves as a stochastic random walk model, as Ljung-Box test null hypothesis of independent distribution could not be rejected for any lag. Then, in the same terms of coordinate analysis, probability of these three specific segments appearing in a non-meiotic validation dataset was assayed as a negative control. Sigmoid fitting coefficients for the LOWESS signal of windowed amplitude transform (Figure 1F) had significantly different distributions across the non-meiotic validation dataset and the positive meiosis experimental data, asserting that this kind of classification into those three specific segments is exclusive for the meiotic prophase (Supplementary Figures S4 & S5). In terms of period and frequency, Power Spectral Density, as well as Complex Morlet wavelet Spectrogram (Figure 1G), display 2.77·10^−3^ Hz or 6 minutes as the dominant frequency or period throughout the first segment (*high nuclear motion*). During the *medium nuclear motion*, the period was significantly reduced up to around 8.3 min. On the other hand, a lack of significant peaks apart from those at very low frequencies are indicative of the noise-like behaviour at the third segment of *low nuclear motion*, agreeing with ARIMA analysis results.

**Figure 1.**
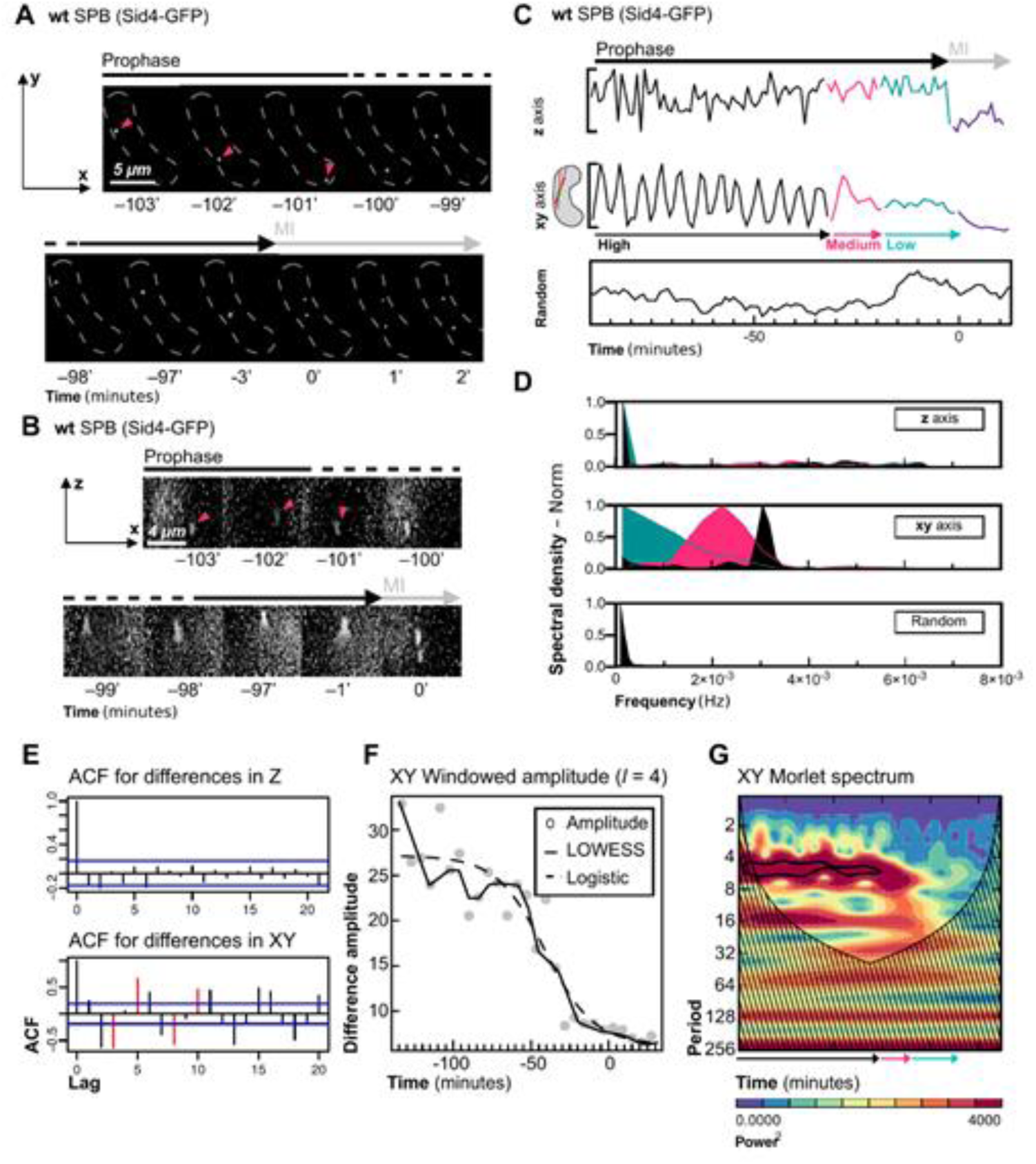
SPB 3D imaging reconstruction and analysis reveals three-segmented meiotic prophase in fission yeast. Selected frames for SPB representation along XY (A) and XZ (B) axes, displayed as the result of fluorescent Sid4-GFP marking. Timepoint 0 is denoted as the beginning of chromosome separation, after SPB duplication. (C) Computed Z, XY projected and simulated Random Walk trajectories with (D) their associated spectral density plots. Proposed and annotated trajectory segmentation is indicated. Colour code for each of the three phases correspond to colours in each of the spectral density area plots, this is, *high nuclear motion* (black), *medium nuclear motion* (magenta) and *low nuclear motion* (blue). Results for each of the statistical tests applied to consider segment significance are shown: (E) autocorrelograms showing significant lags in red, explaining the autoregression model of order *p* AR(*p*) fitting for the whole stages in Z and XY axis, displaying their AR(1) and AR(3) models, respectively. The order means the number of immediately preceding values in the series that predict the value at a given moment. (F) LOWESS local regression, used as the smoothing approach before time-series segmentation, as well as its associated windowed amplitude, with its best-fitting logistic function, whose predicted parameters will be used as the similarity metric among cases. (G) Computed Complex Morlet wavelet plot for the current XY example, with its Power Spectral Density colour code. Black outlines inside the spectrum represent 99.9th percentile significance, usually around 6 to 8 minutes. A total number of 9 cells reaching MI, filmed in three batches, were analysed.

To confirm the new three-stages segmentation from the analysis of the SPB movement, we carried out a complementary approach by the analysis of chromosome movement throughout meiotic prophase (via Hht1-CFP tagging). Moreover, systematic morphological characterization with descriptors, namely circularity, convexity, area and axis length, was assayed throughout meiotic prophase (Figure 2A-B). Consistently, applying the same segmentation protocol in terms of windowed amplitude variance than used for the SPB movement, nuclear morphological information is segmented into the same three phases as in the XY case. During the *high nuclear motion* segment, all morphological descriptors show the most considerable variance across all segments, coinciding with the oscillatory behaviour of the movement. In the *medium nuclear motion* stage, amplitude variance decreases with a similar rate as in SPB trajectories, calling for stabilization of all values throughout the *low nuclear motion* period, with all descriptors varying in values within a normal distribution (Shapiro-Wilk p-value > 0.05) until MI takes place. At this point, circularity, convexity and axis ratio reach their maximum, minor and major axis are minimally different, and size reduces by half with the onset of nuclear divisions (Figure 2C-E). Eventually, negative control validation testing reached a similar result as it did for the trajectory case, sustaining that both trajectory and morphology changes across time are unique to meiotic prophase when compared to non-meiotic nuclei and synthetic morphological descriptors time series (Figure 2F-G).

**Figure 2.**
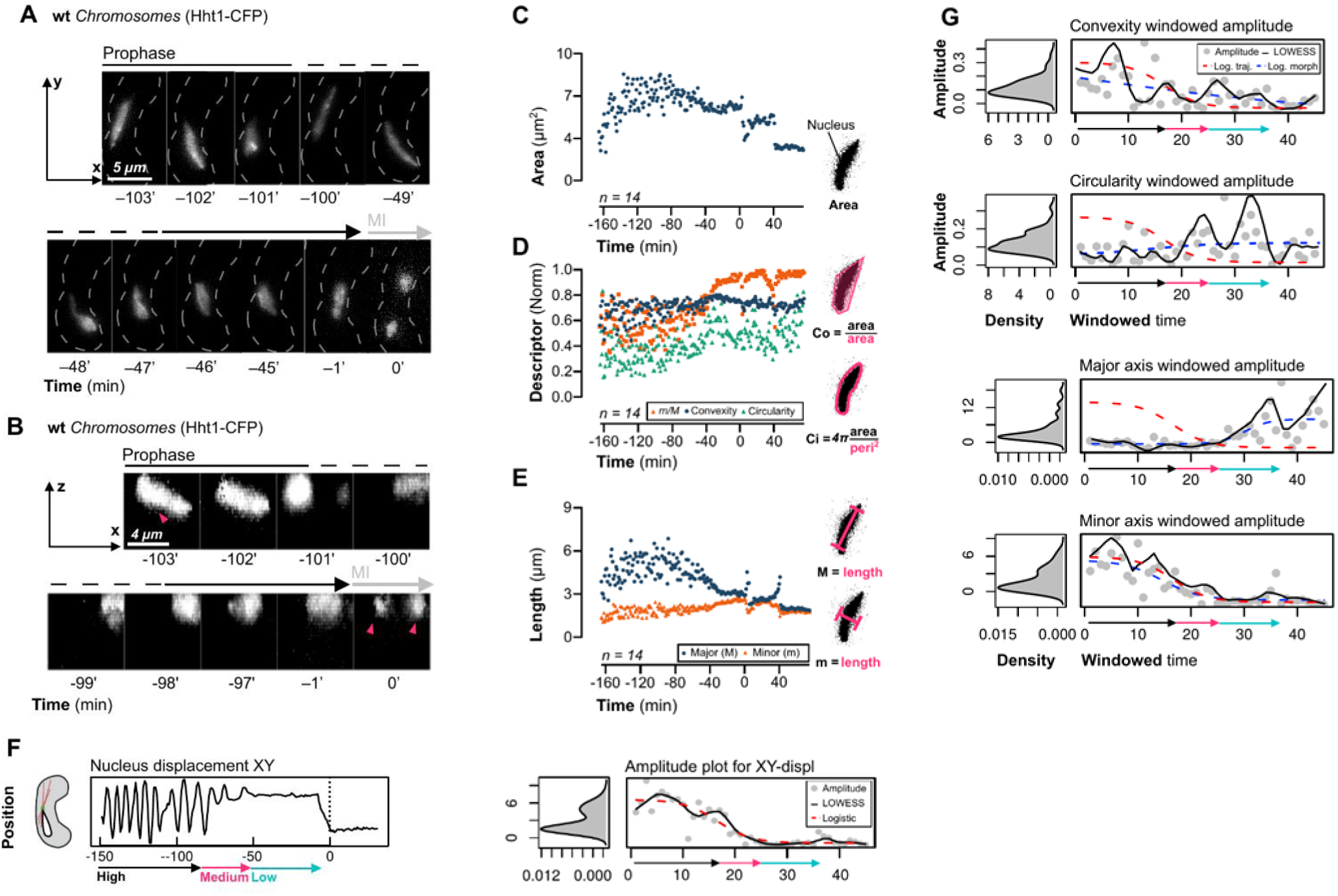
3D fluorescence imaging, reconstruction and analysis of the whole nucleus confirms the existence of three phases during meiotic prophase. Selected frames of cells harbouring the endogenous CFP-tagging of the histone 3 variant, Hht1, are shown along XY (A) and XZ (B) axes; pink arrows mark the nucleus of study. Computed morphological descriptors are shown: (C) Area in microns versus time, (D) normalized convexity, circularity and ratios versus time and (E) Axis length, in terms of major and minor axes in microns, versus time. (F) XY displacements and its associated Windowed amplitude (grey dots), LOWESS (solid black line) as well as the sigmoid fitting (red line) and the significant segmentation calculated, with the previous colour code. (G) Morphological analysis (blue lines and grey dots) in the same previous terms, comparing the segmentation, being derived from the LOWESS-Logistic descriptors of the trajectories (red line). Windowed time is the result of applying windowing to original signals, taking into account the window length (*l* parameter) previously chosen as optimum (Supplementary Figure S2).

### Our analysis reveals new differences in the behaviour of the canonical horsetail mutants

Apart from the more in-depth behavioural characterization of the meiotic nuclear oscillations in a *wt* setting, our interest was to generate a useful tool to identify new horsetail mutants using screening approaches as well as to extract more information from the known nuclear-movement defective mutants. The most common mutations which lead to reduced nuclear oscillations are via deletion of the dynein (Dhc1) and the meiotic-specific microtubule-organizing centre Hrs1. Previous studies have shown slightly more penetrance in *dhc1*Δ nuclear movements defects compared to *hrs1*Δ cells ^15^. In order to establish similarities and differences between both mutants and test our computerized approach, we characterized as previously explained for the *wt* strain, *dhc1*Δ and *hrs1*Δ meiosis in terms of both SPB trajectory and total nuclear displacement-morphology.

Segmentation analysis for the nuclear counterparts revealed, for both *dhc1*Δ and *hrs1*Δ, three Lavielle selected segments: *high, medium* and *low nuclear motions*. Although global amplitudes are lower than in the *wt*, the relative changes in amplitude and frequency during the progression of the time-series still arise. In terms of nuclear morphology, *dhc1*Δ and *hrs1*Δ could be segmented into its matching three phases (Figure 3A). Interestingly, spectral analysis for each segment detected significant differences between the *dhc1*Δ, with no significant 99th percentile values for any frequency, and the *hrs1*Δ mutant, where segment 1 (*high nuclear motion*) displays significant dominant frequencies corresponding to a period of around 8.5 minutes (Figure 3B-C). Moreover, no significant differences in terms of segmentation or spectral densities were found along with *dhc1*Δ strain concerning their SPB-only trajectories counterparts, whereas *hrs1*Δ strain showed spectral incoherencies around the 15 to 9 minutes period. According to previous observations, we noticed a more dramatic reduction in SPB velocity in *dhc1*Δ cells compared to *hrs1*Δ conditions ^15^ phenotype that we were able to quantify in ≈0.16 μm/min and ≈0.65 μm/min, respectively (Figure 3D-E). To sum up, using our approach, we concluded that while *medium* and *low nuclear motions* stages show no significant differences between *dhc1*Δ and *hrs1*Δ strains, *high nuclear motion* segment show a clear difference between both strains. Hence, we think that our new tool will be useful to detect more slight variations in meiotic nuclear motion in a battery of meiotic-defective mutations.

**Figure 3.**
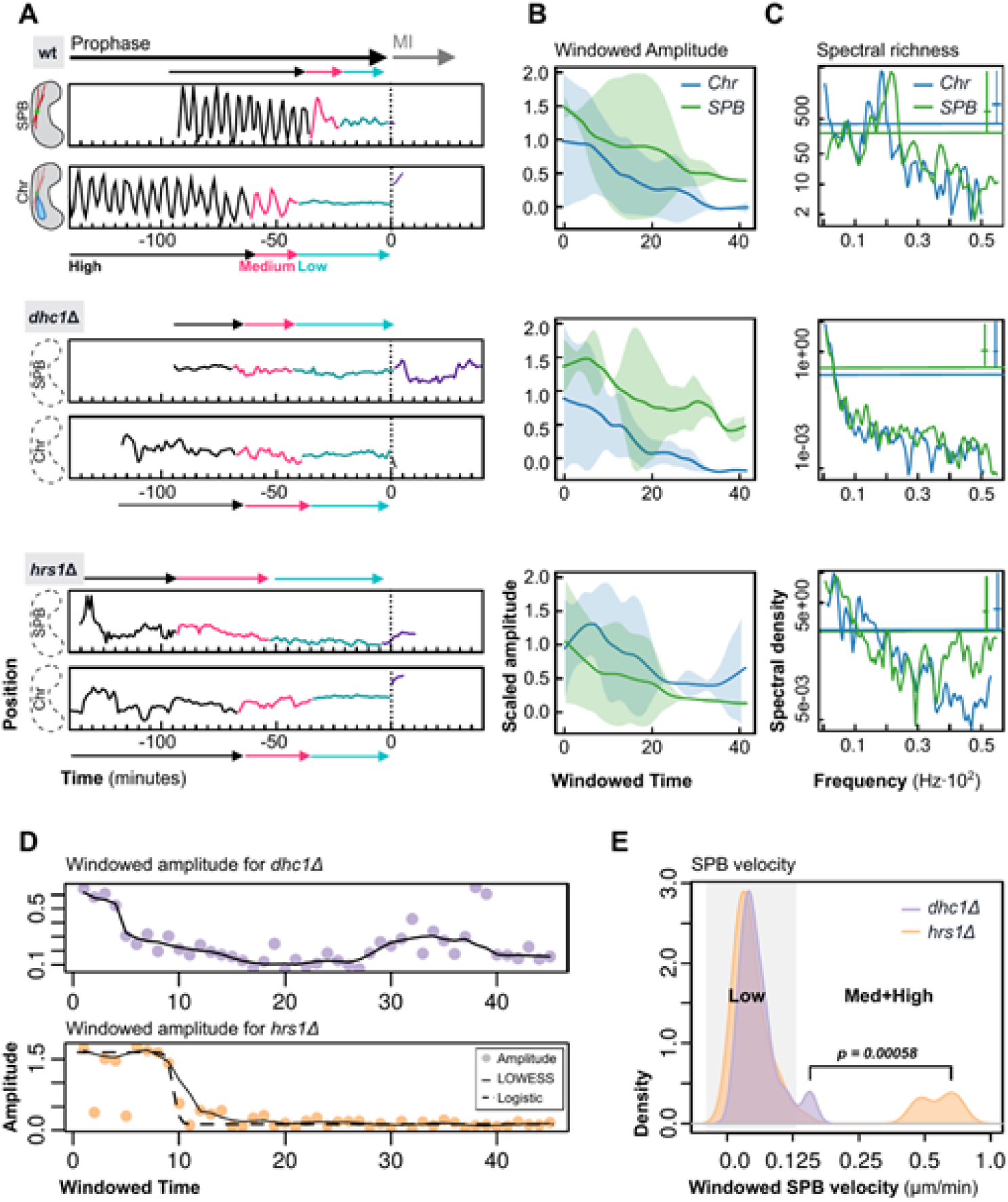
*dhc1*Δ and *hrs1*Δ strains differ in *high nuclear motion* stage but not in the *medium* and *low* segments. Analysis of 3D fluorescence imaging, reconstruction and analysis of the whole nucleus during meiotic prophase in the defective horsetail mutants *dhc1*Δ and *hrs1*Δ marked by Hht1-CFP (chromosomes), and the SPB, marked by Sid4-mCherry. For each listed strain, (A) computed trajectories and significant segmentation are shown for SPB (top) and whole nucleus (bottom) data, with the same previous colour codes (*high, medium* and *low* nuclear motion stages), (B) normalized windowed amplitude plots with range area, as its LOWESS interpretation, for all cells belonging to each strain and (C) its associated global spectral density plots with vertical significance bar indicating the Chi-square distribution 95% CI, and the horizontal bars as the 5th percentile lower bounds. Green colour represents SPB associated data, whereas blue indicates the whole nucleus. In terms of the windowed amplitude, all cases show the first segment with greater amplitude ranges (*high*), ending with small range segment (*low*), both connected by a descending amplitude stage (*medium*). For the whole sequences, significant peaks are those inside the Chi-squared CI. (D) Examples for individual windowed amplitudes, as in (B), for the mutant strains. Total number, as Chr+SPB, of 14+9 (*wt*), 10+9 (*dhc1*Δ) and 12+11 (*hrs1*Δ) cells were analysed. Cells were filmed in at least three independent batches. (E) Density plot for the previous examples, segmented into corresponding *low* and *medium+high*, as there are differences in average SPB velocity across phases; taking into account that x-axis units represent velocity distributions in average microns per minute, with highly significant *t-test* revealing a difference in the means and, thus, in their distributions.

### SPB velocity is not affected by the loss of the telomere bouquet

A key question understanding the molecular mechanisms controlling meiotic nuclear motion is the identification of relevant regulators controlling the SPB movement. It is well-established that microtubules and dynein play a crucial role in nuclear movement dynamics ^7,24^; however, it is a still open question how the chromosomes themselves impact on the SPB movement. The connections between telomeres and the SPB, the telomere bouquet, allow the chromosomes to follow the SPB movement; one possibility might be that the SPB would modify its velocity in cells without the bouquet due to a movement without cargo, although previous data suggesting to discard that hypothesis ^15^ is still debatable ^11^. To establish the patterns of the SPB and nuclear behaviours in cells with and without the bouquet, we carried out the analysis of *bqt1*Δ cells, which are 100% deficient in the telomere bouquet formation ^26^. Interestingly, we found different behaviour when SPB displacement and nuclear counterpart were analysed in *bqt1*Δ settings. SPB trajectory could be divided into the three previously defined stages: *high, medium* and *low motions*, meaning that SPB movement is not changing the velocity by the loss of the bouquet (Figure 4A-C). Significantly, we noticed evidence of displacement behaviour differences, being highly statistically significant, between *bqt1*Δ and *wt* settings, with standard deviations of 2.96 versus 1.78, respectively. Also, windowed amplitudes often reach higher values, although with not enough significance, in *bqt1*Δ cells with respect to the rest. We think that this population of cells represent the cases of the alternative centromere-SPB interaction in the absence of telomere-SPB since centromeres and telomeres have shown interchangeable roles during meiotic prophase in fission yeast ^11^; these transitory attachments then increase the mass carried by the SPB and the motor proteins, causing trajectory alterations. Thus, the sensitivity of our approach can detect the SPB trajectory displacement when SPB and centromeres accidentally interact.

**Figure 4.**
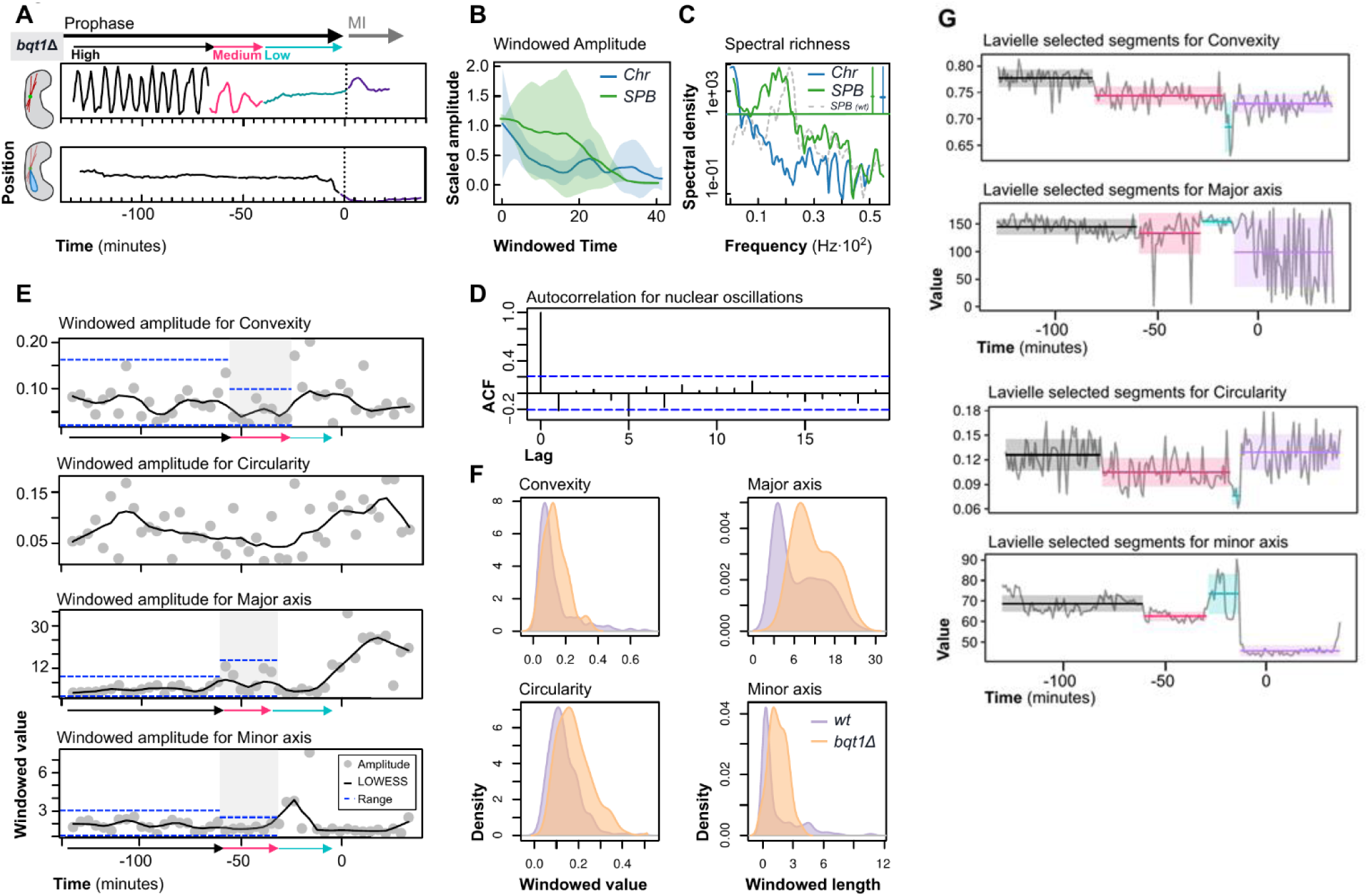
Averaged constant SPB trajectory spectrum and velocity ranges increase in the telomere bouquet defective *bqt1*Δ strain. 3D fluorescence imaging, reconstruction and analysis of the whole nucleus and SPB during meiotic prophase. (A) computed trajectories for SPB (top) and whole nucleus (bottom) cases, as well as significant segmentation, are shown; (B) windowed amplitude plots with range area after LOWESS calculation. (C) Associated global spectral density plots with vertical significance bar indicating 95th percentile Chi-square distribution CI, and the horizontal bar as the 5% lower bound. For the mutant strain, green indicates SPB associated data; blue, nucleus data. The discontinuous grey line is the *wt* SPB spectral density, with comparative purposes. In essence, telomere bouquet loss in the *bqt1*Δ mutant supports the broader significant spectral richness displayed, as a higher variance of SPB velocity due to transitory SPB-centromere interaction (D) autocorrelogram as the ARIMA analysis for the nucleus only motion, with the highest significance at lag 0; (E) windowed amplitudes for each of the morphology descriptors, excluding size, with LOWESS associated Lavielle segmentation, with the same colour code for each stage as before. Blue broken lines represent lower and upper bounds. All descriptors show highly significant segmentations, except circularity, which is normally distributed, as the nucleus is highly static in this mutant, lacking perceivable circularity variation. (F) Densities for each of these descriptors are shown, both in the *wt* (purple) and *bqt1*Δ (orange) cases, (G) Lavielle segmentation for the original data, with no differences calculated nor LOWESS description, in terms of nuclear morphology; colour code is maintained from the *high, medium* and *low* segmentation, as well as MI (purple). Similar segments to those in panel E are conserved along with axis length descriptors; also, convexity and circularity LOWESS-segmentations are contained inside the global, non-modified original signal segmentation. Total number, as Chr+SPB, of 9+12 *bqt1*Δ cells, filmed in at least three independent experiments, were analysed.

On the other hand, analysis of chromosome dynamics showed that nuclear displacement was not significantly different from negative controls, displaying mostly Brownian like behaviour (Figure 4D). In fact, *bqt1*Δ cells meiotic prophase could be divided, in terms of morphology, into three significant stages, corresponding with the free and attached nucleus and SPB, as a requirement for first meiotic division concurring to the *medium* and *low nuclear motion* stages. Within the context of this strain, some filmed cells show instantaneous nuclear morphologies which resemble those of the *hrs1*Δ, as having low amplitudes but significant frequencies and morphology-trajectory correlation at the upper 99th percentile (Figure 4E-G).

### Nuclear morphologies and SPB trajectories are divided into four different motifs

To further characterise the molecular bases of meiotic nuclear oscillations, we investigated, in more detail, the behaviour of nucleus and SPB in order to extrapolate possible uncharacterized patterns of motion. Remarkably, we detected some significantly single segments displaying internal bimodal distributions (Supplementary Figure S5). Hence, these distributions may exist since trajectory and morphology stages are made up of smaller motifs, or due to random noise distribution because of the filming process or other factors outside the cell. To this end, a matrix-profile based algorithm applied across all trajectory data against a validation set revealed four significant motifs which we defined as *motif A, B, C* and *D* (Figure 5A), not in the negative validation set. Those were mostly detected on both *high nuclear motion* and *medium nuclear motion* phases (Figure 5B). Amplitudes and frequencies of infinite time series built up from each motif subset are significantly different and with homogeneous peaks (Figure 5C-E). Regarding its internal development, *motif A* has a sine-like appearance with a predominating 6 minutes period, and *motif B* shows mostly at the ending points of *high nuclear motion* stage, as the transition between itself and the *medium nuclear motion. Motif C* has dual periodicity, as in terms of trajectory it accounts for a brief returning to the departure point after a directional change, then regular oscillation restores. In this aspect, this will be referred to as trajectory anomalies, in the sense that it is substantially different from the traditionally studied *motif A* in terms of frequency. *Motif D* has a displacement shape characteristic of the *medium nuclear motion* towards the cell centre. With all the information obtained, the relative abundance of each motif in phases 1 and 2 was calculated (Figure 5F).

**Figure 5.**
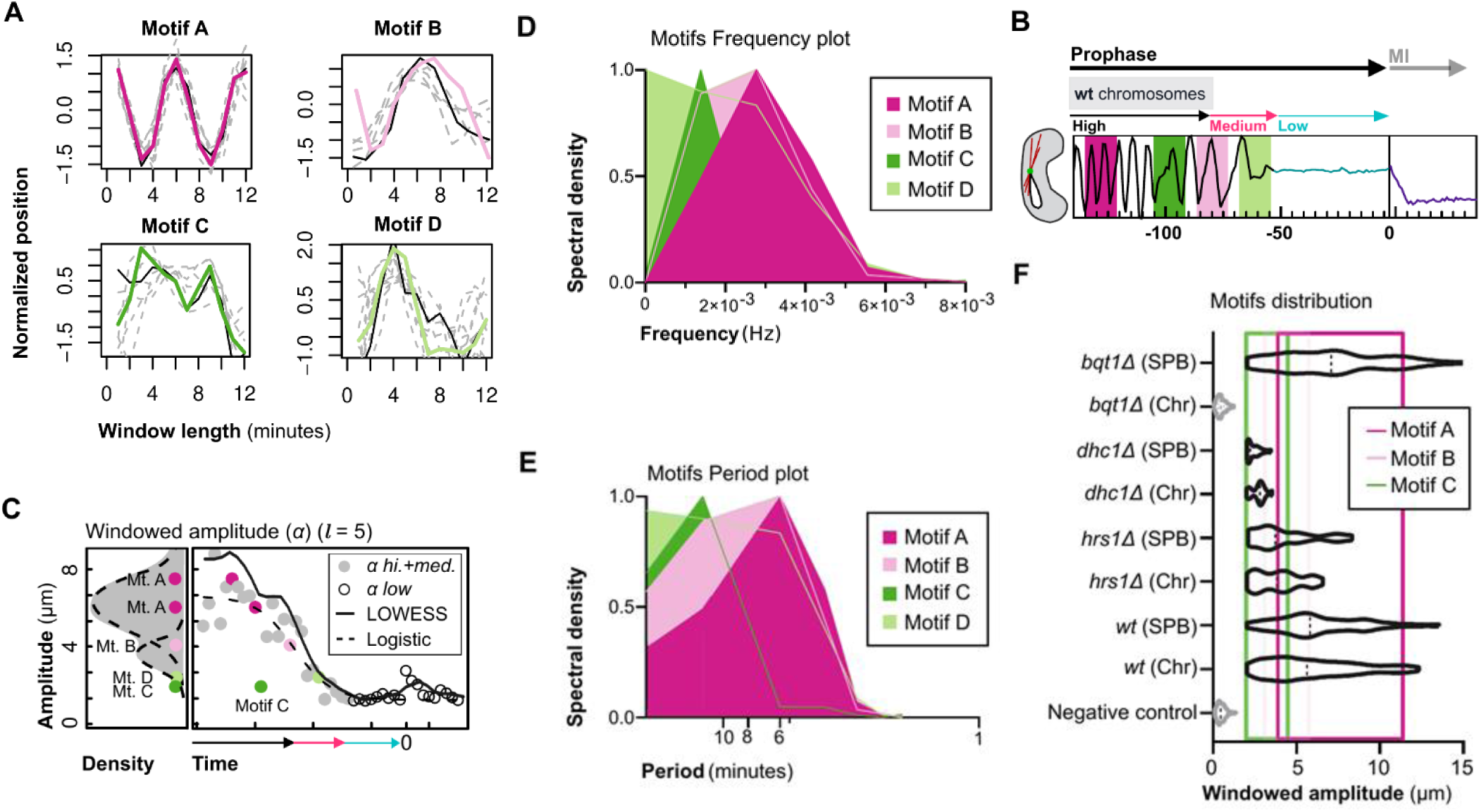
Displacement motifs discovery for the whole explored dataset. (A) displays examples and alignments for each of the significantly discovered motifs. (B) represents motifs across a typical segmented horsetail trajectory sample, indicated as the colour outlines for the boxes around small portions of the sequence. (C) Windowed amplitude for the previous example, with its associated LOWESS and Sigmoid-logistic fittings, and the densities with broken lines representing inner normal distributions of the stages *high* and *medium* (grey dots), excluding the *low* stage (empty dots), rich in low frequencies. Colour dots are examples of motif appearance during meiotic prophase, with their projections in the density plot, illustrating the contribution to spectral richness. (D, E) contain the tapered significant frequencies and periods for time series constructed uniquely by motifs. (F) contains data for strain and filming condition motif relative abundance in *high* and *medium* stages. Area of the violin plot occupying each colour, corresponding to each strain and condition, indicates the relative amount of the motif, as each has a different place of dominant Windowed amplitude.

In terms of morphology, descriptor information (Supplementary Table 2) for all meiosis-positive nuclear time series was subjected to FNMM-EM clustering (see Methods), thus obtaining four morphological classes (*i, ii, iii* and *iv* in Figure 6A). During analysis, only four of the five descriptors were considered significant, excluding size, as final classification showed no difference when comparing both scenarios, thus simplifying visualization and later analysis. Classes *i* and *iii* have both a high relation of major to minor axis, low circularities and low convexities, whereas classes *ii* and *iv* have an almost equal major and minor axis, high circularities and high convexities (Figure 6B-C). To sum up, our results reveal an extra layer of complexity behind of meiotic nuclear motion whose biological significance we tried to disclose.

**Figure 6.**
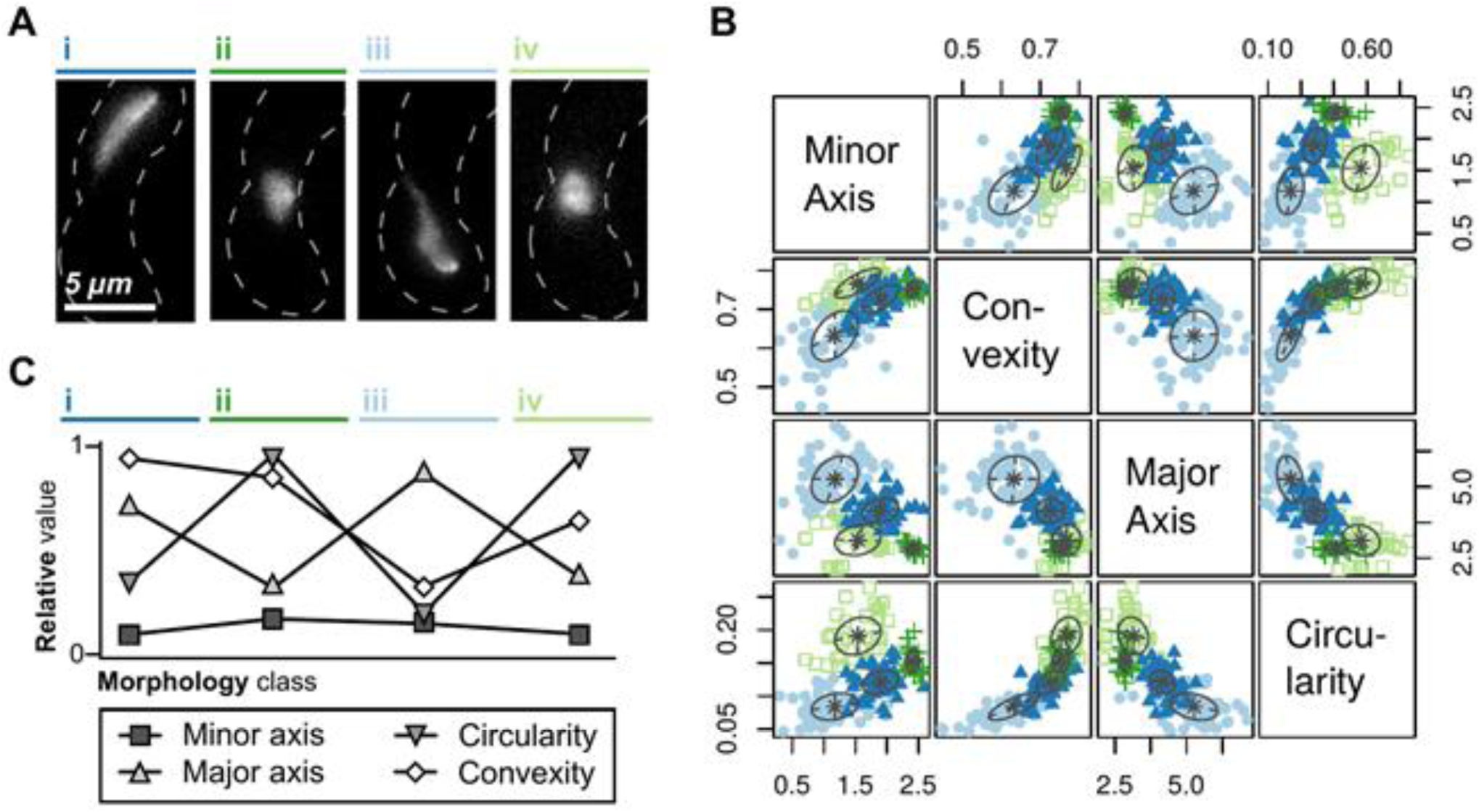
Nuclear morphology throughout meiotic prophase can be divided into four classes. Meiotic nuclear morphology assay for *wt, dhc1*Δ, *hrs1*Δ and *bqt1*Δ strains harbouring Hht1-CFP as in Figure 2A-B.; (A) Representative examples of each morphology class (*i, ii, iii* and *iv*), with the matching legend colour. (B) Paired correlation tests for all chosen morphological descriptors, convexity, circularity, major and minor axis, across all frames and strains, showing different colours and radial outlines for each of the classes detected and their distributions. (C) Summarized graphical representation of features leading to class differentiation, in terms of circularity, convexity and axis length.

### Nuclear trajectory anomalies are linked to morphology abnormalities

Once trajectory and morphological information is segmented and divided into motifs, the possibility of causality relations among them was explored, in order to test whether there are relations explaining morphology and trajectory implications that drive, for each stage within meiotic prophase, different alterations and, eventually, distributional motifs. Observation of the convexity descriptor distribution across strains (Figure 7A) shows a possible relation between motif appearance and lower convexities, especially in the case of motif C, as the morphological motif distribution has a similar shape to that of the trajectory motifs in the previous section (Figure 5F). For each strain, Granger causality analysis was performed (Figure 7B). In terms of possible causation, Granger’s test is widely used in several scientific fields ^27,28^, explaining whether one time series forecasts another and asserting how two functions are related, as one being lagged to the other. Taking the previously calculated amplitude window length optimum as the reference value, Granger test gave significant results (p-value < 0.001) after fitting differenced trajectory data and convexity information, chosen as the parameter best-describing morphology irregularities. In this case, significance means that the time series of convexity data contains enough information that explains variation in the trajectory data, not as an immediate, zero-lagged response but with some delay. Hence, it is worth noting that, as convexity reaches values out of the fluctuation scope, the trajectory has a late response in the form of oscillatory interruptions, mainly, low convexity values associated with morphological class *iii* are likely to be behind the Granger-causation of motif C appearance. As control scenarios, the inverse relationship is the null hypothesis of convexity being Granger caused by morphology, and could not be rejected, since no significant p-values were obtained. Also, data of cells with non-ascus appearance did not support significant Granger causality relations, indicating the uniqueness of these relations during meiotic prophase. Overall, trajectories (*B*) are Granger caused by morphology (*µ*) changes and not contrarily, as represented by the relation *B = µ(t)*. Additionally, the correlation was significant between morphological descriptors compared to convexity, with the maximum cross-correlation value at lag 0 (Figure 7C). Importantly, we concluded that there is a causality relationship between alterations in convexity and trajectory anomalies.

**Figure 7.**
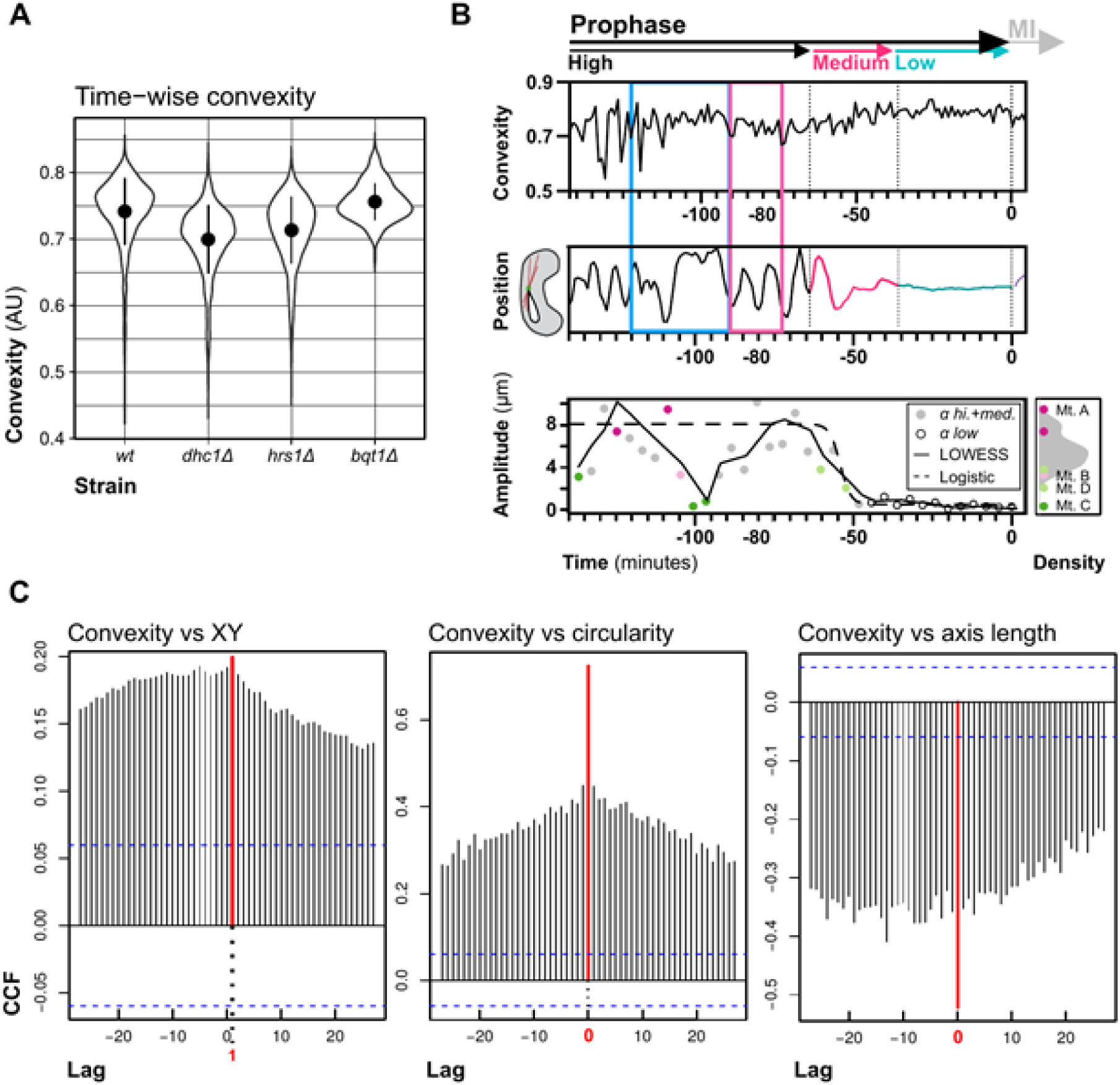
Morphology and Trajectory follow a causal relationship. Morphology-Trajectory causality analysis for *wt, dhc1*Δ, *hrs1*Δ and *bqt1*Δ settings, (A) Distribution of convexity as a highly informative morphological descriptor across time. *wt, dhc1*Δ and *hrs1*Δ show similar distributions, with a high range of convexities indicative of morphological variation across time, with the *horsetail* mutant strains having, in average, lower values possibly linked to abnormal trajectories. *bqt1*Δ case shows less variance in convexity due to highly reduced nuclear motion, although being left-skewed, suggesting morphology variation across time, especially in the convexity descriptor. (B) *wt* example of time-series alignment, in terms of original, non-differenced data for convexity and XY position, with its associated windowed amplitude plot with LOWESS-Logistic fitting and densities excluding the *low* stage, as well as examples of different motifs. As a graphical summary, it illustrates how Granger causality test will be performed, with results with p < 0.05 meaning that there is a Granger-causality relation between sets analysed. A blue box remarks convexity (upper time-series) to motif C (lower) appearance window, and a pink box encloses convexities related to motif A. In (C), cross-correlation function (CCF) plots for the convexity versus XY position, circularity and axis length. Red lines indicate the highest absolute value for the CCF. Blue dashed lines indicate the significance limits, giving a clue from a graphical perspective of positive results in the Granger test.

### Persistent DNA damage leads to nuclear convexity changes throughout the horsetail movement

Granger causality analysis revealed that there are morphological implications, as within the convexity descriptor, affecting trajectory. Next, we wanted to establish which phenomenon is bringing those nuclear morphological alterations, keeping in mind the possibility of these alterations being due to external factors, such as genotoxicity, eventually causing cellular responses. Specially, we will scope motif C appearance, as being linked to convexity changes which can be potentially coupled with molecular genetic alterations. Previous observations have shown that persistent DNA damage extends the telomere bouquet stage once SPB movement is slowed down ^25^. One possibility would be that DNA damage induced during meiotic recombination might not only affect the extension of the bouquet after the horsetail movement, also, alter the chromosome dynamics and morphology during *high nuclear motion*. To test that hypothesis, we analysed an ATP-dependent helicase mutant (*rdh54*Δ) strain which showed a more prolonged post-horsetail stage due to defects in repairing of meiotic DNA double-strand breaks ^25,29–31^. Applying the same methodology, we identified the three segments corresponding to the meiotic prophase: *high, medium* and *low nuclear motion* in both SPB trajectory and nuclear morphology data (Figure 8A-B and Supplementary Figure S6 & S7). Consistently to previous observations, we noticed a longer extension of *low nuclear motion* stage in *rdh54*Δ cells compared to *wt* settings (Supplementary Figure S8, Supplementary Tables 3 & 4). Regarding global motifs, there were no additions nor subtractions to the previously presented pool, as all of them remained the same. Also, Granger causality was significant (p-value < 0.05), suggesting that there is a causal relation between convexity and altered morphology in this case (Figure 8C). In terms of relative abundance of motifs and spectral analysis, both SPB and whole nucleus filming of the *rdh54*Δ strain yielded indistinguishable time-series, as the same distribution was observed for motif C appearance, and both significant period peaks at around 6 and 9 minutes were conserved with almost equal Power Spectral Densities (PSD) (Figure 8D-E). Hence, we found that the motif C is present in all *rdh54*Δ settings strongly suggesting that the presence of this motif might be associated to promote the repairing of double-strand break.

**Figure 8.**
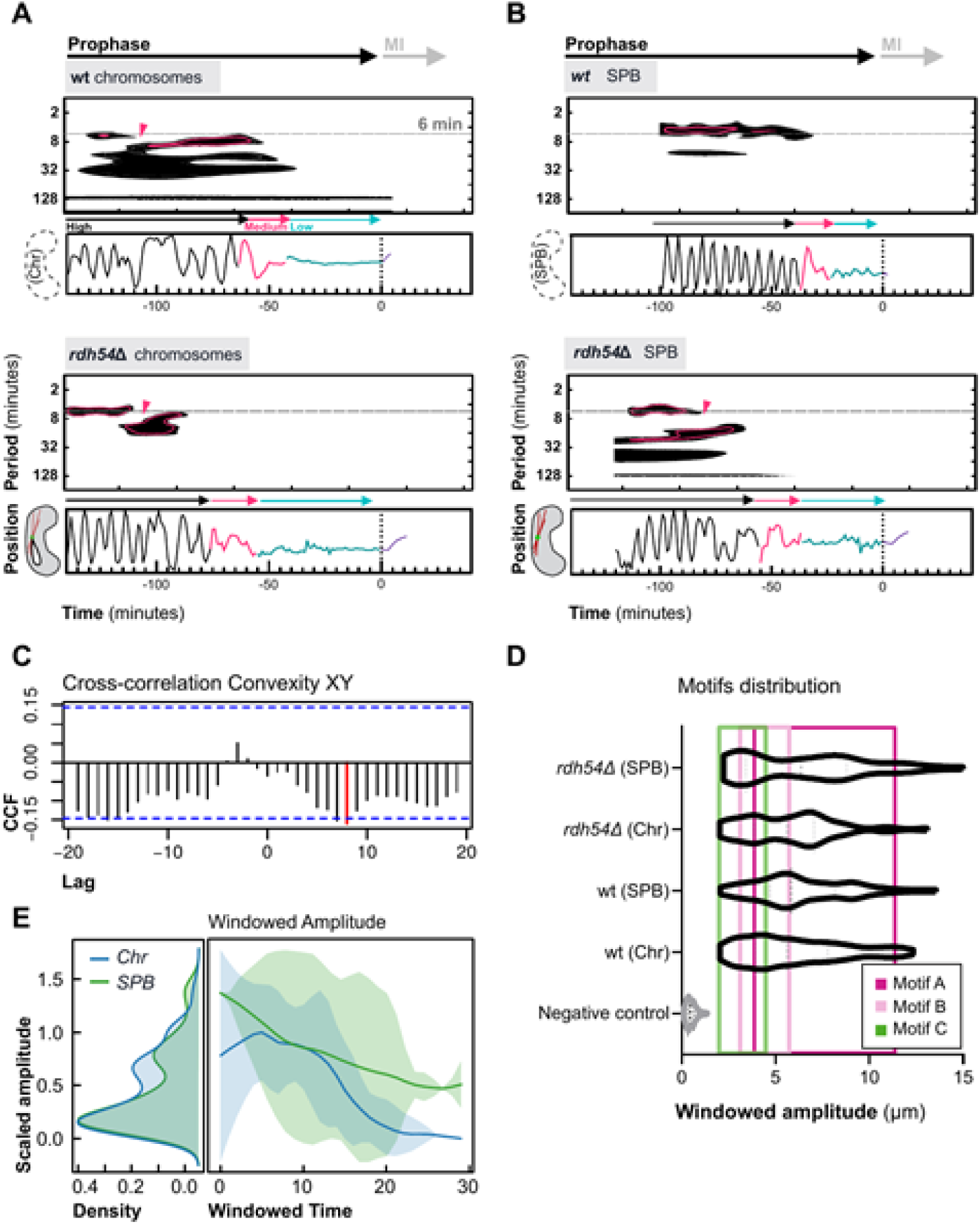
The appearance of Motif C is enriched in *rdh54*Δ mutants. (A-B) Comparison of Complex Morlet Wavelet transformed spectrogram (top) and trajectory time-series data with annotated segmentation (bottom) for *rdh54*Δ and *wt* examples, by whole nucleus (A) or SPB (B) analysis. Pink outlines inside the spectrogram indicate the 99.9th percentile significance, with pink arrows showing points where the leading frequency is interrupted, and dashed grey lines, 6 minutes period. PSD over 90th percentile are black, with the rest being white. (C) cross-correlation analysis for the convexity versus XY position, with a significant lag equal to 8 (red line); blue dashed lines represent significance thresholds. (D) shows relative motif appearance as the area inside a particular colour - uniquely associated with a motif -. Similarities of both SPB and whole nucleus cases for this mutant is disclosed in (E), with the density plots for the three stages, as well as its associated LOWESS fitting with bounds. For these analyses, a total number of 9 (Chr) and 11 (SPB) *rdh54*Δ cells were used. Data was collected in at least three independent batches.

## DISCUSSION

Until now, meiotic prophase in fission yeast has been divided into two phases; (*i*) the *horsetail* stage, characterized by vigorous nuclear movement, where chromosome pairing and recombination take place, the end of this phase has been established by the reduction of intense movement triggered by the degradation of the protein Hrs1, a meiotic-specific MTOC at the SPB ^32^; (*ii*) the *post-horsetail* stage, where persistent DNA damage is repaired, whose end is controlled by the disassembly of the telomere bouquet at MI onset ^25^. These subdivisions are based on the study of nuclear motion stages as being delimited by biochemical milestones. Here, we designed a systematic study founded on the sequential nature of this mechanism, uncovering evidence from their own time and frequency domains. With this aim, we established that meiotic prophase could be segmented into three different nuclear motion stages depending on the *high, medium* and *low* nature of their windowed amplitudes or spectral richness. Moreover, as previous studies regarding this phenomenon have not taken into account the three-dimensional nature of nuclear displacement, our imaging and analysis protocol included a Z-dimension analysis enabling tracking in that direction. In this aspect, it was shown that Z trajectories fitted best with a Random walk model; hence, astral microtubule projections directing the trajectory of oscillation are likely to be random in this plane, as no single type of pattern is prevalent over time ^33^, behaving as an stochastic system. Following with spectral analysis, as frequencies vary, *high* and *medium* stages show systematically non-stochastic behaviours, with strong significant periodicity. Conversely, *low nuclear motion* is for the first time annotated as significantly following a stochastic model, thus being possibly due to SPB and nuclear thermal oscillations of the whole molecular complexes supporting nuclear environment across this period, as zero net displacement is observed ^34^.

In order to establish which are the essentials after the spectral contributions for each of the three stages, mutants annotated as having reduced horsetail movements were analysed; as a result, it was shown that both *dhc1*Δ and *hrs1*Δ backgrounds are easily distinguishable, in terms of SPB velocity and spectral richness. In that sense, it is known that Hrs1 regulates the onset of horsetail oscillations ^19,26^, while Dhc1 allows the oscillation itself acting as the main motor protein ^7,16–18^. Hence, the lack of the motor is producing a more severe phenotype, in which spectral components typical of the horsetail oscillations are lost, as the nuclear structure as a whole is not tightly attached to microtubules which, due to polymerization dynamics reaching cell poles, deliver strong periodicity. On the other side, loss of *hrs1* did not yield this severity, as some residual and significant oscillations were found; when Hrs1 is lacking, Alp4 recruitment to the SPB is diminished but not completely abolished, as it remains dispersed around the nucleus ^26^. Hence, spontaneous interactions may induce subtle but significant oscillations as nuclear structures remain attached to microtubules. Likewise, as nuclear motion is reduced in the *dhc1*Δ case compared to *hrs1*Δ, *low motion* stage duration is inversely proportional, allowing to hypothesize that reduced spectral richness, and consequently nuclear displacement, lead to requiring a higher amount of time during stationary phases right before the onset of MI, substantiated by the requirements of proper chromosome pairing and recombination. SPB velocity, although complying this relation in the *dhc1*Δ and *hrs1*Δ cases, does show more like *wt* values in *bqt1*Δ, being this variable discarded in that previous statement. Moreover, nuclear morphology variation yields an almost equal segmentation to motion counterparts, as notably seen in *bqt1*Δ: linkage of both SPB velocity and morphology alterations, particularly convexity, are partially sustained by transient SPB-centromere interaction ^11^ which elicits brief trajectory responses. Though, extensive nuclear morphological oscillations happening in the *medium* phase compared to the *low* phase when segmenting convexity and axis length remain unexplained, as transient interactions do not happen when the SPB is at cell poles; consequently, inner signals from the own nucleus, such as nucleoproteins molecular integrity, may be relevant in the self-regulation of oscillations.

Additionally, the classical *horsetail* model cannot explain how each segment has different inner distributions. These microvariations can be approached as four motifs in terms of trajectory and morphology. Discarding that this segmentation is due to random events, we wanted to regard their individual biological meanings: *rdh54*Δ strain has been described, among others, as having a more extended *post-horsetail* phase ^25^ and altered nuclear morphology during meiosis and reduced recombination of markers ^30^, being a right candidate for assaying intrinsic DNA damage caused by a reduced pace of repairing ^29^. Compared to its *wt* counterparts, there is a similar behaviour in terms of trajectory segmentation, observable motifs and their distributions, except for the case of SPB-only filming. As longer wavelengths were operated to excite GFP or mCherry in marked Sid4 SPB component, it is known that in those conditions genotoxicity is minimal, in contrast to conditions where shorter wavelengths, hence with more energetic fluorescence beams, are used to excite CFP marked histones, with literature data supporting that assumed higher genotoxicity is significant ^35^. As revealed, extended duration of the *low nuclear motion* stage, corresponding to the previously described *post-horsetail* phase, is related to increased accumulation of DNA damage. Also, motif C relative appearance in the SPB-only filming of the *rdh54*Δ strain is higher from what was observed in the *wt* case, being *low* stage duration also significantly greater. Therefore, there is a tight correlation between aberrant trajectories and DNA damage accumulation, both presumably caused by impaired DNA repairing or by external genotoxic factors. Also, it may be hypothesized that nuclear structural alterations detected as low convexity during meiotic prophase and distributional linked to motif C, as observed in morphological class *iii*, can be linked to DNA damage accumulation since a Granger causality between convexity and trajectory has been demonstrated. Wherefore, motif C is the outcome of a structural alteration that interrupts the oscillatory pattern, probably deriving into an increase of tension forces between the nucleus with respect to the SPB and attachment complexes; in that case, their summation leads to slowing down and returning, with a contribution of viscous drag forces within chromosomes ^36^. As described in the literature, this form of morphological disorders can be caused by defects in chromatin cohesion ^37,38^. Taking a global perspective, the model proposed in this work refines previous models explaining nuclear oscillations during meiosis prophase, as it considers a wider time window and spectral richness, arising three behavioural segments with an even more complex internal development composed of motifs. Hence, chromosome movement is not only being carried by astral microtubule dynamics, motor and coiled-coil protein localization as a form of extrinsic regulation, but is also linked to intrinsic mechanisms ultimately mediated by nuclear morphology in virtue of molecular highlights such as DNA damage.

To sum up, we show that meiotic prophase takes place in three different stages: *high, medium* and *low nuclear motion*. It was observed for the first time that only XY-bidirectional displacement gives oscillatory-like information, as Z dimension is dependent on the XY axis, following a Brownian like behaviour by itself. It was also demonstrated that *high* and *medium* stages are divided altogether into four motifs, in terms of morphology and trajectory. These motifs had different appearance rates across mutants and wavelengths used during microscopy; especially, motif C occurrence, as instantaneous interruptions of the primary oscillations, and its correlation with a more concave than normal looking nucleus morphology. Our study revealed that those trajectory anomalies correlate with cellular events, specifically, those related to DNA damage accumulation, possibly triggering morphology disorders. Hence, these can explain how the nucleus, as an intrinsic factor itself, regulates the oscillatory patterns apart from the extrinsic cytoplasmic motors and structures. Consequently, this opens the possibility of this methodology for characterizing nuclear alterations during meiotic prophase as indirect markers of DNA damage accumulation and its associated cellular responses, opening its relevance up to the applicability while studying alterations during gametogenesis in higher eukaryotes, such as humans.

## METHODS

### Strains used and growth conditions

All strains used throughout this study are specified in Supplementary Table 1. Cells were maintained in standard growth conditions on YES-agar media in Petri dishes at 32°C ^39^. Stocks were maintained in YES-Glycerol media at -80°C.

### Meiosis induction and sample preparation

Homothallic (*h90*) haploid strains were grown on YES medium plates at 32 °C. Then, sufficient biomass was plated on a SPA agar plate, then waiting up to 6 hours at 28°C. Biomass patches were checked for a sufficient amount of mating cells under the light microscope. 50 µL of 0.2 mg/ml soybean lectin (Sigma Aldrich) were put in the centre of a 35 mm glass culture micro-dish microscopy (Ibidi Gmbh) and let sit for 2 minutes. Biomass was taken with a sterile toothpick and re-suspended in 200 µL MilliQ and, after lectin was recovered from the plate and let dry, 100 µL of the cell suspension was put in the centre of the plate for 4 minutes. Eventually, EMM minimal medium was used to perform successive washes of the remaining biomass and to fill the plate with up to 3 mL.

### Fluorescence microscopy

Time-lapse image data was obtained using a DeltaVision widefield microscope system (Applied Precision, Issaquah, WA), equipped with a Photometrics CCD CoolSnap HQ camera, a UV filter and an Environmental Chamber set at a constant temperature of 28°C, checked to be stable enough 30 minutes before filming. Images were taken with a 100x 1.4 NA oil immersion objective, every minute for 3 hours over 20 z-planes at a 0.4 µm step size. Hence, pixel size is 0.06 µm, giving a total amount of 181 temporal points per cell recorded. Values for exposure time and transmittance were 100 ms/32% respectively, for fluorescence channels, and 50 ms/50% for the DIC channel. An Argon lamp and specific sets of filters were used to excite different markers; 436/10 nm for CFP marked Histone Hht1, and 470/40 nm or 492/18, respectively, for GFP or mCherry marked SPB component Sid4.

### Initial image processing

Raw *dv* extension files generated during live-fluorescence microscopy image acquisition were processed using a custom binary file wrapper written in Python with libraries PIL, OpenCV2 ^40^ and struct. Maximum slice intensity or global sum z-projections were applied to the fluorescence channels of the image stack depending on the signal to noise (S/N) ratio, followed by Otsu’s thresholding, thus obtaining binary images. Cells with *ascus* appearance were annotated as meiotic positive with a Convolutional Neural Network, based in Pytorch ^41^, previously trained with a dataset of 150/150 positive/negative examples, and a 70/30 proportion for both training and validation sets. Also, depending on the specific excited marker, Chromosomes or SPBs were detected and treated as <u>b</u>inary large <u>ob</u>jects (blobs), as the group of connected pixels in a binary image. Blobs resulted from 255-positive pixel 8-connectivity of the binary image after blurring with a Gaussian filter, across all frames. Accordingly, blobs inside positive meiotic ROIs were marked as positive samples, as the rest of negative meiotic samples were used to correct frame drifting and bleaching.

### Time series data reconstruction

Blobs detected in the fluorescence channel at the initial image processing stage were calculated their underlying morphological descriptors, namely, circularity (*c*), major axis (*θ*_*M*_), minor axis (*θ*_*m*_), convexity (*µ*) and total area (*A*), as defined in ^42^. Aspect ratio is calculated as the axis ratio *θ*_*m*_ /*θ*_*M*_. A set, *P*, includes globally detected and linked particles, *p*, which is individually defined as a tuple

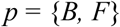

where *F* is defined as the global set of descriptors in time, *f*

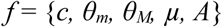

and where *B* is the trajectory defined as the time series

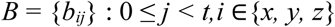

being *j* the time point, *t* the total time in minutes, and *i* the three-dimensional Euclidean space, as *x, y* or *z*. Particles were reconstructed from individual blobs in each frame by their positions and morphological descriptors as input data. For that purpose, the *trackpy* Python library was used ^43^, with a maximum null linkage of 4 frames, a maximum travelled distance of 100 pixels and a minimum appearance in 80% of frames. An output struct contained, for each particle, timepoints and their related positions and morphological descriptors.

### Time series analysis

Particle location in time is 0-1 normalized to cell size and its position. Previously derived time-wise individual particle information was initially reduced in terms of dimensionality, considering trajectory and morphology separately. Trajectory was also separately analysed in two steps, one involving the differenced form *D* derived from the *x* and *y* directions, as

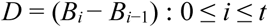

and other involving the *z*-direction with respect to time, respectively. Trajectory windowed amplitude α of the differenced form *D* was calculated to perform time-series segmentation. It is defined as the discrete function

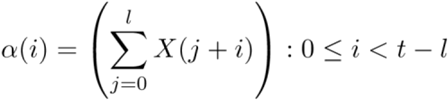

where *l* is the minus-one optimal window length in time units, *X* the position function for the current particle *p* with respect to *x, y* or *z* directions. For subsequent segmentation analysis, <u>L</u>ocally <u>w</u>eighted <u>s</u>catterplot <u>s</u>moothing (LOWESS) regression was applied as a Savitzky–Golay smoothing for α(i). A sigmoid curve fit was also performed ^44^. As for the general form of this sigmoid σ(x) function, with parameters *b, c, d* and *e*, with *x* as the α(i) windowed amplitudes:

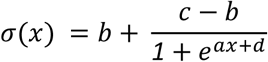

For time series segmentation, the *segclust2d* R package was used ^45^. Optimal segmentation is associated with a likelihood value; hence, chosen segmentation is calculated as having the Lavielle optimally selected likelihood ^46^. Hence, a trajectory is redefined as the joining of several segments, as the tuple

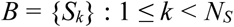

where *S* is the segment *k*, which is enumerated from 1 to the least segment, *N*_*s*_. Accordingly, segments *S* acquire the previous trajectory definition, as

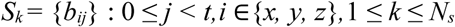

For each segment *S*, preliminary ARIMA analysis was performed in order to determine whether Brownian motion, as a case of random walk time series, is the predominant underlying stochastic process driving nuclear dynamics ^47^. Random walk *W*_*N*_(t) with steps δ = 1/n is defined by induction as

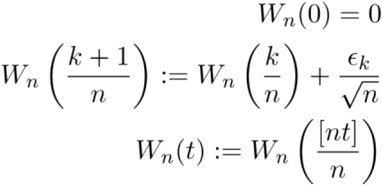

with the integer part of *a*, and identically independently distributed random variables

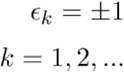

Hence, as a stochastic process, the Auto-regressive model of order *p* AR(*p*) is defined as

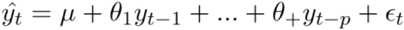

allowing a random walk process to be modelled as an AR defined process with *p* = 1 and θ > 0

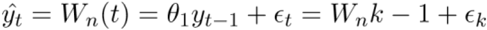

Random walk positively fitting segments are discarded in subsequent steps. Positive fitting of an ARIMA model is defined by minimum BIC and AIC values, p-values of coefficients being < 0.05, normal distribution observed in qq Plots, and significant Ljung-Box test during residuals analysis, as the differenced form of the applied time series. For segments with non-stochastic trajectories, the spectrum is calculated ^48,49^. Morlet wavelet transform is applied considering a sampling rate of 1 minute or 60 seconds, equal to 0.01667 Hz. Morlet spectrograms were represented with a 0.999 significance level to consider dominant frequencies. All dominant peaks were stored as components of a feature vector for each segment in each particle. Individual strain feature vectors are joined in terms of spectral density and frequency, applying a Gaussian Mixture Model (GMM) with Expectation-Maximization (EM) algorithm to discover significant global peaks ^50^. If no segmentation was initially possible or no apparent differences are detected within segments, a trajectory is homogeneous, and ARIMA analysis is equally applied to it.

### Validation sets construction

Trajectory and morphology data from both blobs annotated as meiotic negative and synthetic signals were used as negative controls for the meiotic prophase biological process. On the one hand, time series reconstruction protocol for negative meiotic blobs followed the same steps as in the positive experiments. Signal generation was performed with a custom function written in Python, as a white noise random signal, with equality across all the Power Spectral Densities, as well as random walk simple stochastic processes as defined before. These data were equally treated, in terms of class structure, regarding their positive meiosis counterparts.

### Motif discovery and significance analysis

Trajectory motifs for each of the non-stochastically driven segments are described as *m*, after Matrix Profile for each input segment is calculated with the *tsmp* package by using the STOMP algorithm, with Dynamic Time Warping as the distance metric and Hierarchical Clustering as the clustering method. Morphology descriptors are classified using a Finite Normal Mixture Modelling (FNMM) model with EM algorithm across axis, convexity and circularity descriptors at once, as performed within the *Mclust* package ^51^. Granger causality approach is used as the method described in ^52,53^ to consider correlation or causality relations between sets of time-series data, attempting to determine whether one series is likely to influence change in the other. Hence, a series change is modelled through lagging another series, being Ω and π these models, with *k* as the number of lags, eventually described as

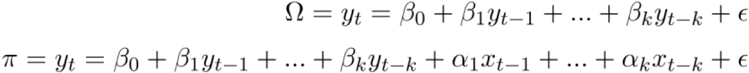

where residual sums of squared errors (SSE) are compared by a standard Wald test.

### Statistical testing, figure plotting and computational resources

Correlation analysis of morphology and displacement motifs, linear and curve fitting, Shapiro-Wilk tests for the assay of normal distribution within spectral peaks and segment duration data as well as scatterplots, cluster visualization, colour spectrograms, qq Plots, density plots and boxplots were generated in R with *astsa, Mclust, dplR* and *ggplot2* packages. Figures were assembled using Inkscape. Code for the workflow and sample data is publicly available in the repository https://github.com/danilexn/Tronos

## ACKNOWLEDGEMENTS

We thank Pedro A. San-Segundo, Andrés Clemente-Blanco and Antonio J. Pérez-Pulido for critical comments on the manuscript; Kazunori Tomita for the *rdh54*Δ strain; Alejandra Cano for technical support; and the CABD microscopy facility technician Katherina García and Alejandro Campoy for their helpful advice. We would like to thank the Genetics Department and Springboard lab for their useful discussion and comments, especially Víctor Carranco for technical support. This work was supported by Spanish Government, Plan Nacional PGC2018-098118-A-I00 and Ramon y Cajal program, RyC-2016-19659 to AF-A; and by the Spanish Education and Professional Formation Ministry, Research Collaboration Grant to DL-P The CABD is an institution funded by Pablo de Olavide University, Consejo Superior de Investigaciones Científicas (CSIC) and Junta de Andalucía.

## AUTHOR CONTRIBUTIONS

DL-P. and AF-A. designed the study; DL-P. performed all experiments; AF-A. acquired funding and supervised the project; DL-P and AF-A. wrote the paper.

