## Supplementary Figures for "Identification of a meiosis-specific chromosome movement pattern induced by persistent DNA damage"

### SUPPLEMENTARY MATERIAL

#### *Supplementary Figures*

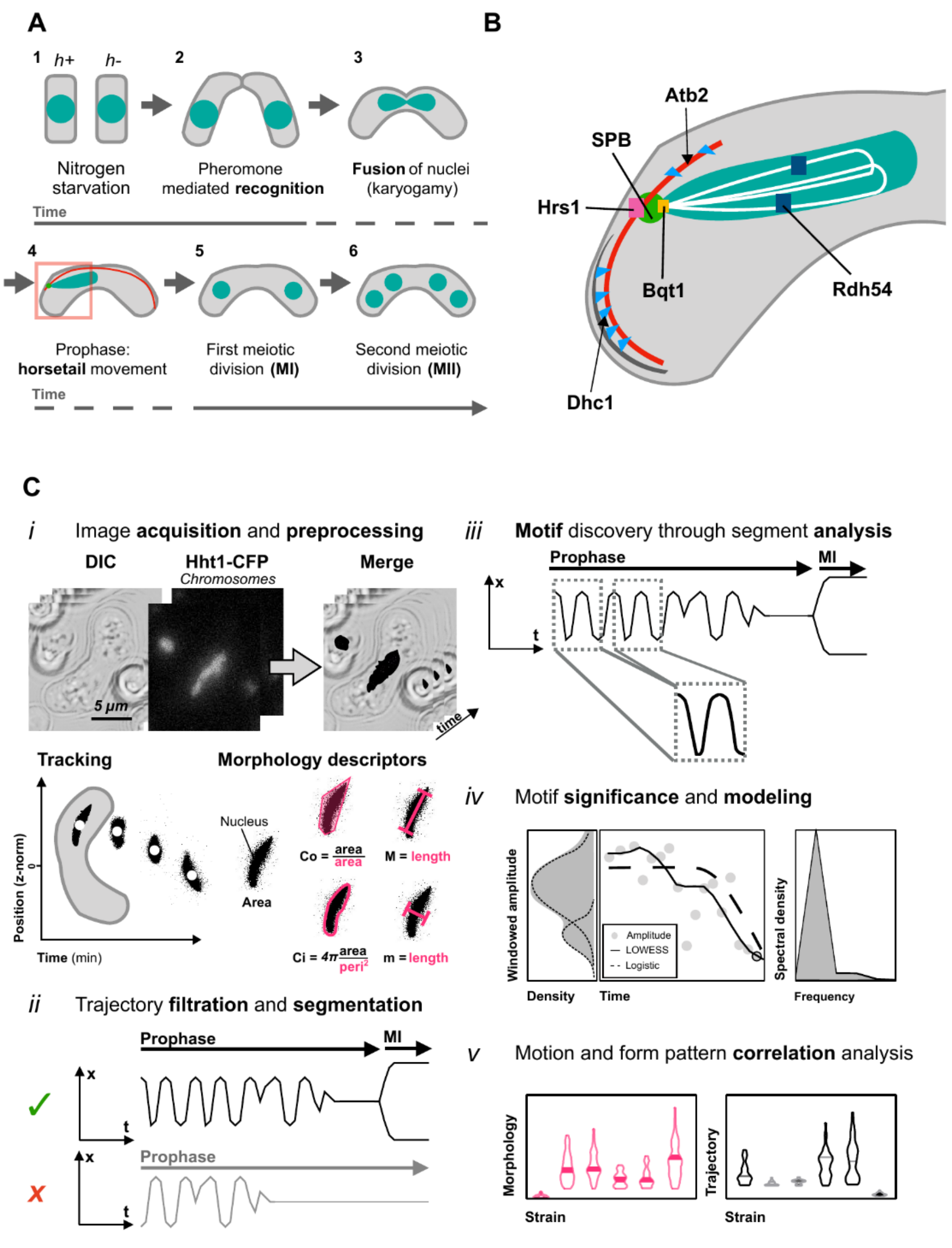


**Supplementary Figure 1**. **Processes occurring across meiotic prophase and workflow summary**. (A) concerning relative time; $h^+$ and $h^-$ haploid cells, in a nitrogen-starvation medium (1), tend to (2) secrete hormones that allow their mating, after which (3) nuclei from both cells fuse in a process called karyogamy; (4) consequently, horsetail oscillations begin, until (5, 6) two consecutive rounds of segregation take place (MI and MII), leading to spore formation. (B) Schematic representation of biological structures involved in the horsetail nuclear oscillations, naming essential structures considered throughout this study. (C) Sequential order as followed during work-flow execution: (i) as the first stage of image processing, blob extraction and data reconstruction, in terms of trajectory and morphology, (ii) only data from cells yielding MI is considered positive for subsequent analysis; (iii) motifs are discovered through a matrix-profile algorithm in joined segments from the whole dataset. Classification into categories is applied to morphological descriptors; (iv) motifs are studied for each significant distribution, or segment, in terms of its belonging; (v) correlation between all kinds of descriptors is studied, in terms of distribution similarity and cross-correlation.

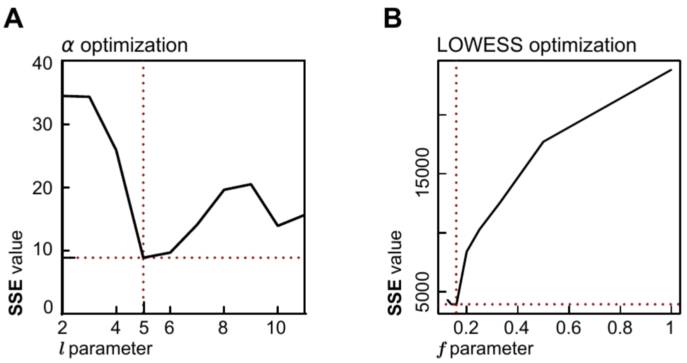


**Supplementary Figure 2. Optimal parameters fitting for analytical descriptors, for the parameters used in Windowed Amplitudes in Figure 1;** considering optimality when the sum of squared errors (SSE) is minimum across the own signal difference and windowed amplitude or LOWESS fitting, (A) Window length ($l$) calculation for Windowed amplitude descriptor, revealing an optimal at $l = 5$. Once this value is obtained as a departure point, for the calculation of windowed amplitudes, l = 4 achieves the best resolution, allowing to assert small variations which l = 5 neglects. (B) is the smoothing span calculation for LOWESS, with an optimal chosen value of 0.16 or 1/6.

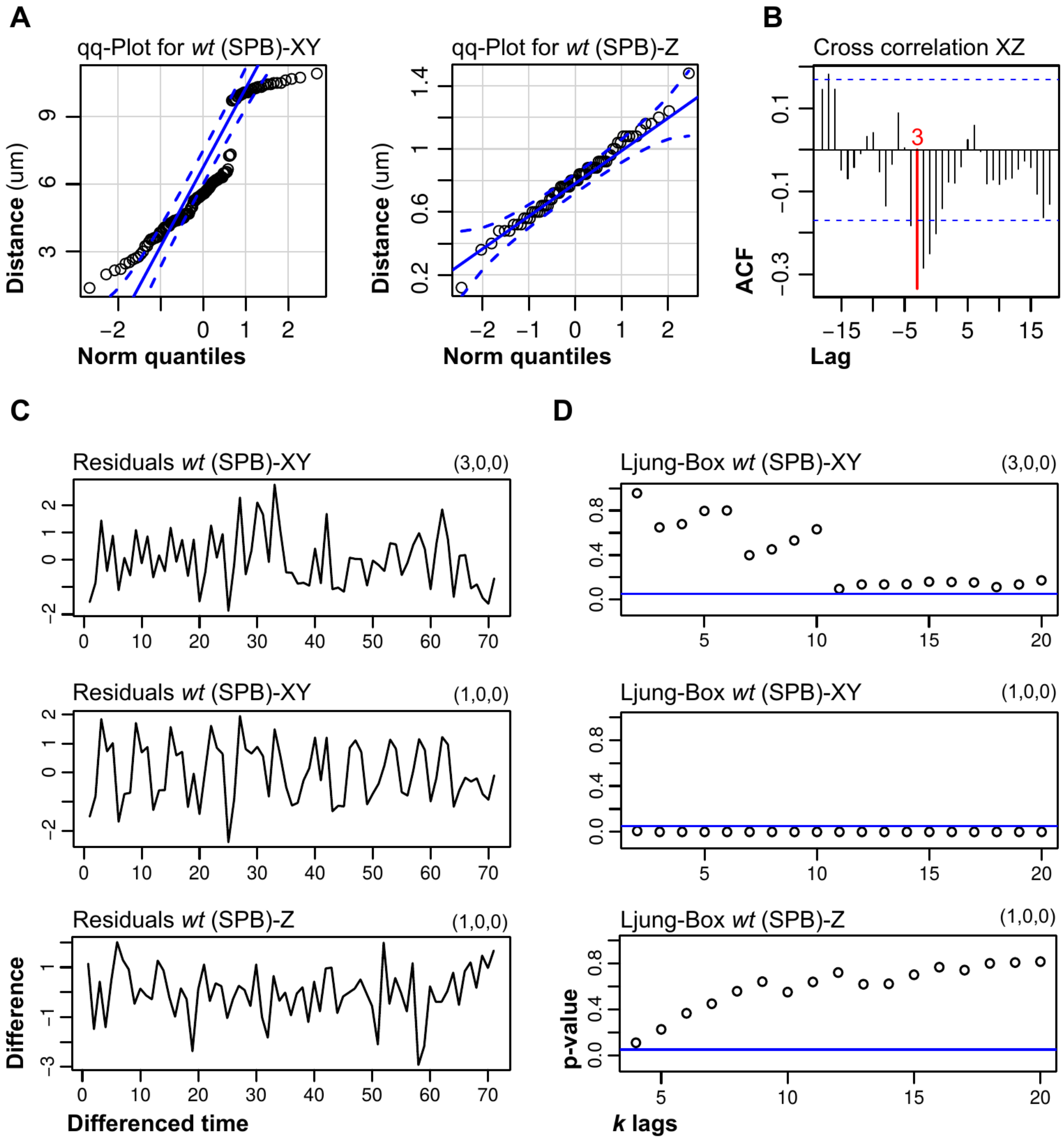


**Supplementary Figure 3. ARIMA analysis for *wt* strains across XY and Z axis in Figure 1.** In order to reveal (p, d, q) as the standard AR, difference and MA orders to give a clue of motion stationarity or strong periodicity at higher lags different from zero. ARIMA orders allow the description of a response variable, in terms of the autoregressive, integration and moving-average models, that is, how the influence of previous values explains present values. (A) normality assay through qq-Plot visualization for XY and Z trajectories, showing that the motion across Z-axis is normally distributed, thus suggesting Brownian like displacement. (B) cross-correlation between X and Z is accounted to correct for possible cell position effects that may give Z periodicity. For that purpose, differenced forms of both types of trajectory are subjected to automatic ARIMA order calculation; following the parsimony principle, (C) coefficients are calculated as shown in the top right of the residuals plot for each case. (D) Ljung-Box tests reveal the significance of the Brownian-like behaviour of the Z-axis, as p-values are above the significance threshold (0.05) for all lags. For the XY case, an AR process of order 3 is most likely to be the driving process than an AR(1), as shown by best Ljung-Box test significances and more normally distributed residuals.

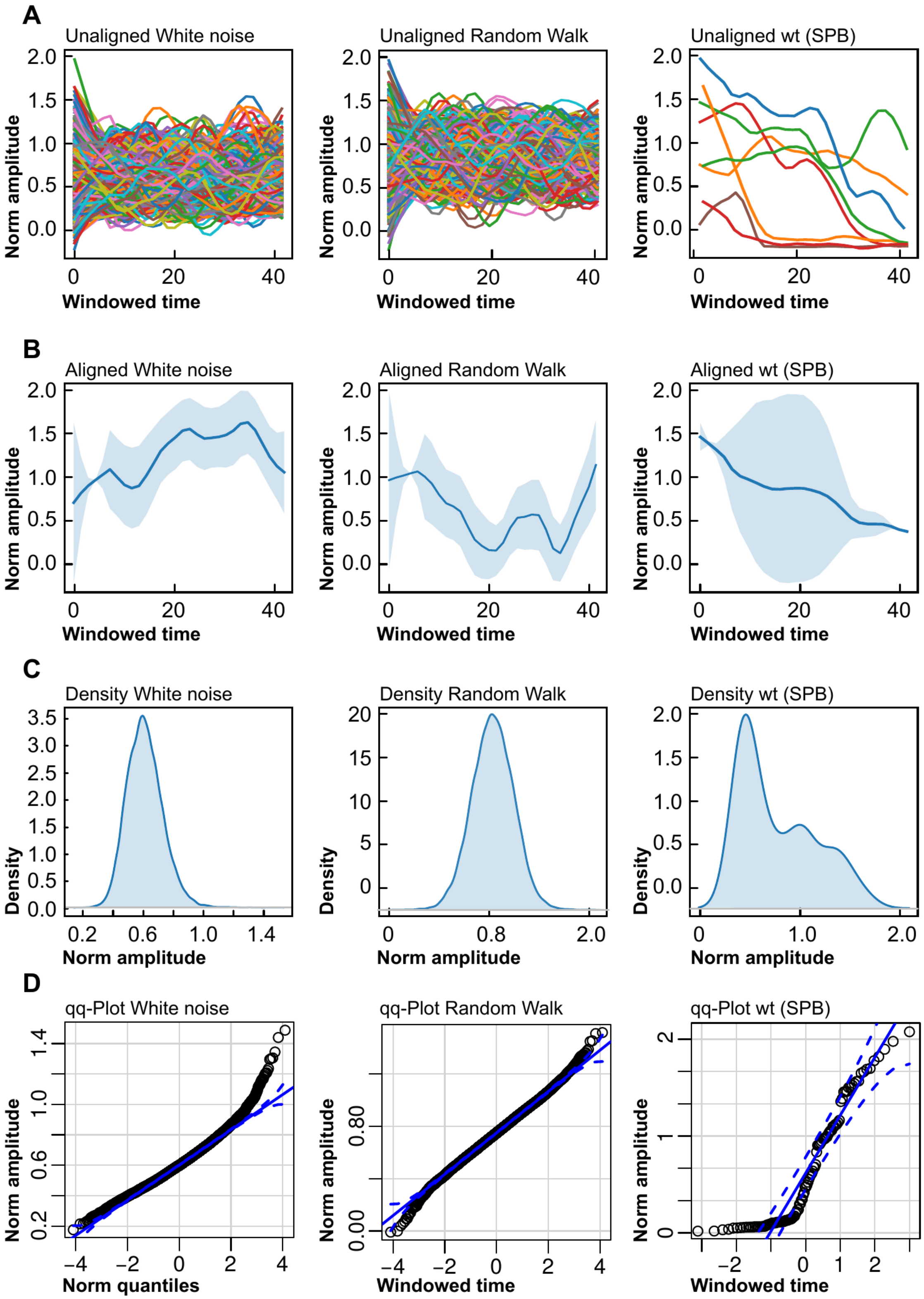


**Supplementary Figure 4. Negative control set comparison with respect to *wt*, as shown in Figures 1 and 2, for the whole dataset.** From top to bottom, (A) unaligned and normalized windowed amplitude plots for signals, both synthetic and experimental, (B) same previous data, aligned and with range areas, (C) distribution for the alignments of windowed amplitudes, suggesting the differences in the underlying models, as well as the qq-Plot graphically stating normality across samples (D).

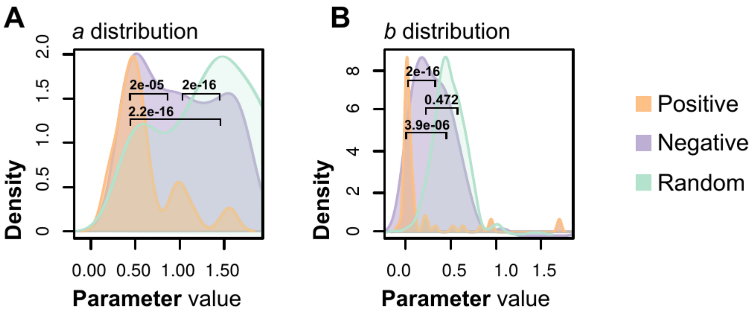


**Supplementary Figure 5. Sigmoid fitting for positive and negative controls;** histograms for the distributions of the coefficients *a* and *b* in a typical sigmoid-logistic function (see Methods), for the meiotic positive, meiotic negative and random walk trajectory data. Welch test p-values for each pair, considered significant under the 0.05 threshold, as well as legends for each dataset, are shown.

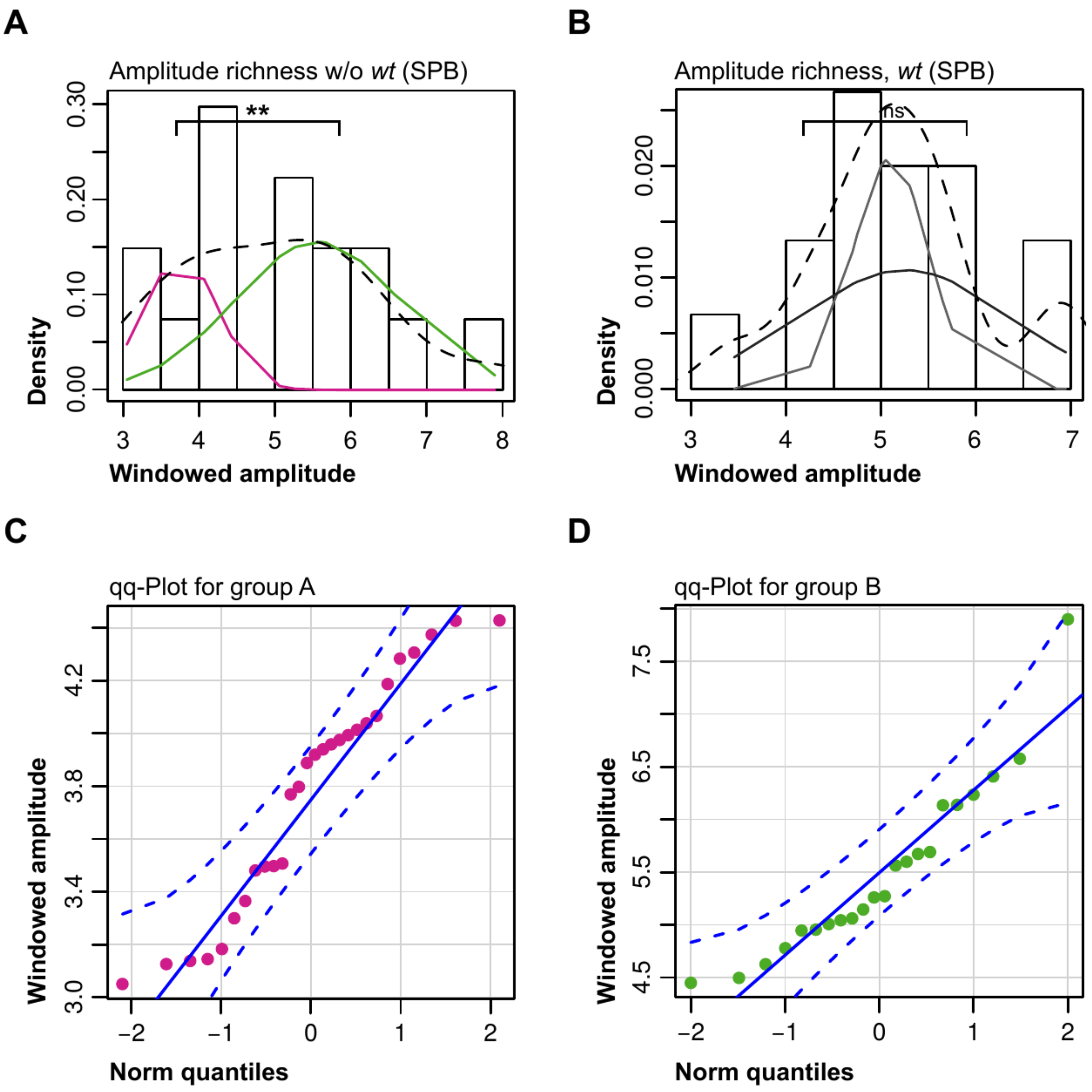


**Supplementary Figure 6**. **Windowed amplitude distribution of segments *high* and *medium* in *wt* and *horsetail* impaired as the cases shown in Figure 3**. Shapiro-Wilk normality test for each individual segment revealed that, except for the *wt* (SPB) case, all-time series data, including morphology and trajectory, did not correspond to normal distributions. Proper GMM-EM fitting of the global windowed amplitudes distribution revealed for *horsetail* impaired mutants (A) and the *wt* case (B) how bimodal distribution is likely to apply, with significantly different distributions only in the first case (Welch *test* p-values are shown), each colour represents the different underlying normal distributions; Shapiro-Wilk tests and qq-Plots (C, D) yield and show significant results for the two distributions disclosed in the case of the mutants. P-value legend: (**) < 0.01.

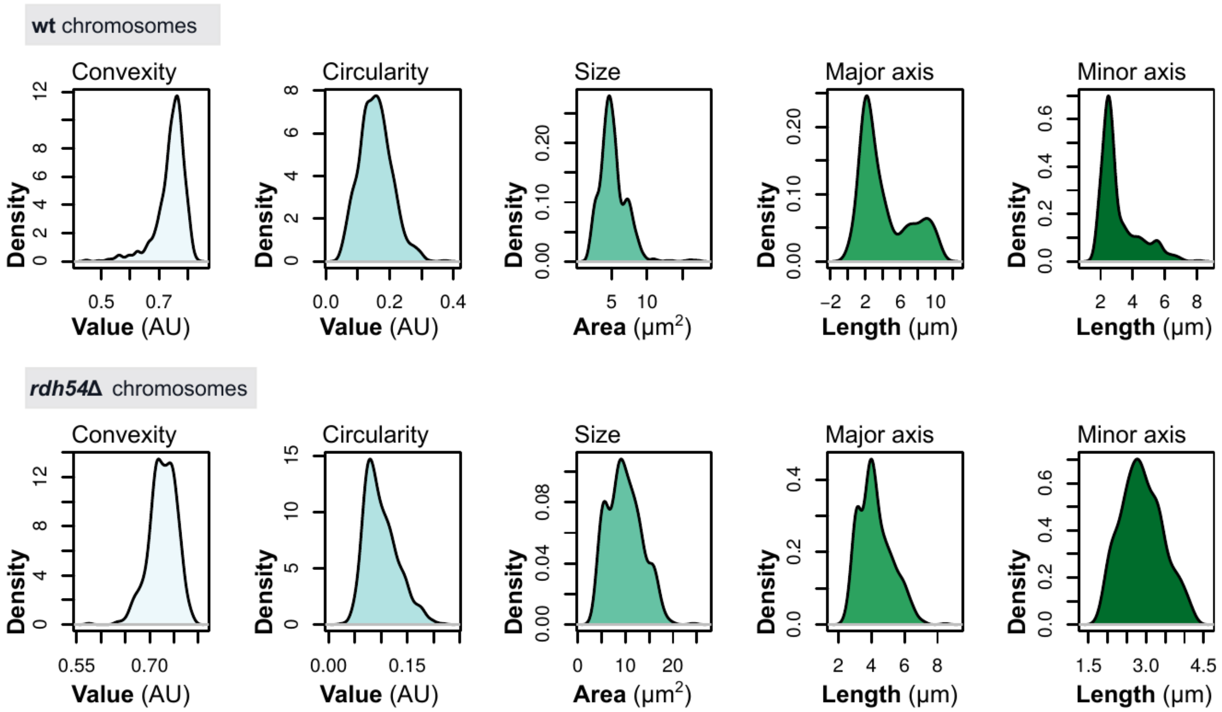


**Supplementary Figure 7. Morphology descriptors distribution for *wt* and *rdh54*Δ regarding Figure 8.** All five descriptors are shown, often having multimodal distributions indicative of possible time-series segmentation about these. Most notably varying descriptors, from one strain to the other, are convexity and axis length. Nuclear size is the most coherent, whereas circularity is more skewed in the *rdh54*Δ, probably being due to frequent smaller convexity values.

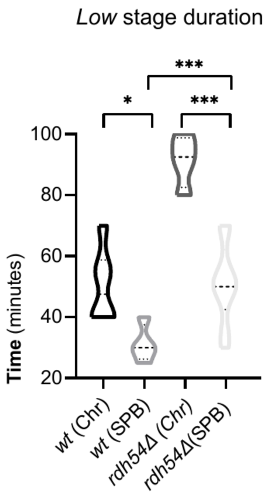


**Supplementary Figure 8. Time length within the *low nuclear motion* stage**. Violin plot represents data for the *wt* and *rdh54*Δ cases, for both Chromosomes and SPB filming data. Significant differences were detected by One-way ANOVA test for each strain when comparing their SPB and whole nucleus counterparts. P-value legend: (*) < 0.05, (***) < 0.001.

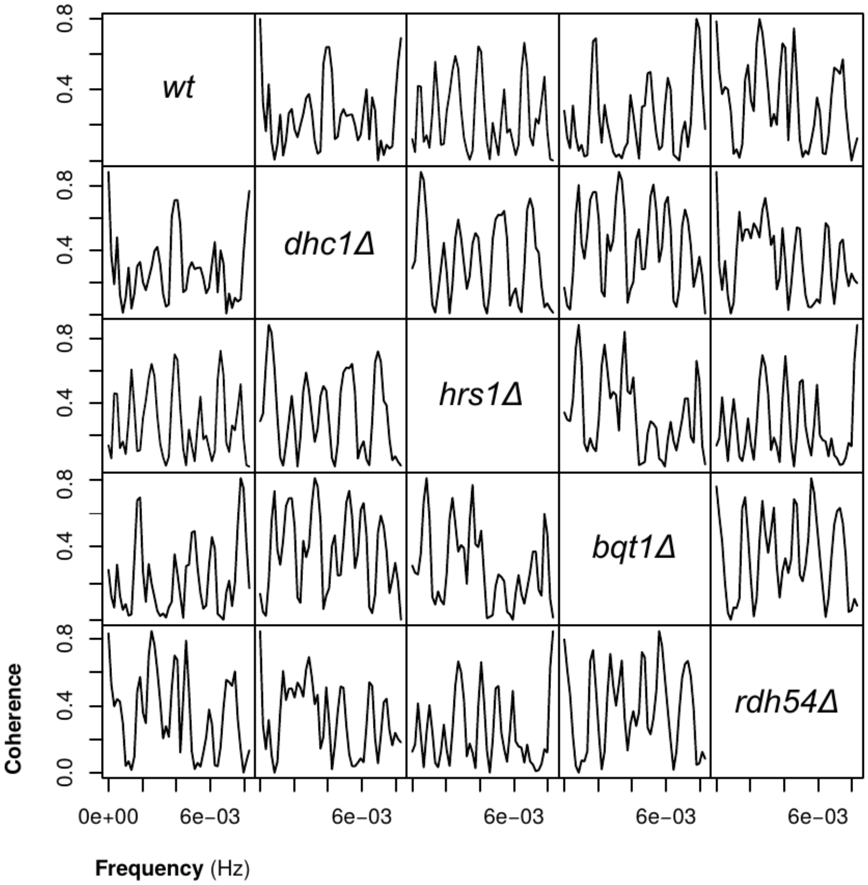


**Supplementary Figure 9. Spectral coherence for all strains analysed throughout this study.** Higher peaks represent highly conserved frequencies across, being this case the total nuclear motion. Highest global coherence similarities, calculated as the area under the curve, are as follows: *wt* with *rdh54*Δ, *dhc1*Δ with *bqt1*Δ and *hrs1*Δ with *dhc1*Δ.

*Supplementary Tables*

**Supplementary Table 1. Strains used throughout this study;** specifying lab strain collection identifier, genotype and origin.

| Strain | Feature | Genotype | Origin | Ref. |
|---|---|---|---|---|
| *AFA226* | *wt* | h90 his3-D1 Pnda3-mCherry-Atb2:aur1 Sid4-GFP:KanMX6 Hht1-CFP-his | Lab stock | -- |
| *AFA824* | *hrs1Δ* | h90 ade6-M210 his3-D1 leu1.32 Hht1-CFP:his3 Pnda3-mCherry-Atb2:aur1 Sid4-mCherry:NatMX6 mis6-GFP:KanMX6 hrs1::hygMX6 | JCF10361 | (Fennell et al., 2015) |
| *AFA825* | *dhc1Δ* | h90 ade6-M210 his3-D1 leu1.32 Hht1-CFP:his3 Pnda3-mCherry-Atb2:aur1 Sid4-mCherry:NatMX6 mis6-GFP:KanMX6 dhc1::hygMX6 | JCF10378 | (Fennell et al., 2015) |
| *AFA826* | *rdh54Δ* | h90 ura4-D18 taz1-YFP:KanMX6 Hht1-Cerulean:ura4 Sid4-mCherry:NatMX6 mei4-mCherry:NatMX6 rdh54::zeoCV | KT2475 | (Moiseeva et al., 2017) |
| *AFA827* | *bqt1Δ* | h90 ade6-M210 his3-D1 leu1.32 Hht1-CFP:his3 Pnda3-mCherry-Atb2:aur1 Sid4-mCherry:NatMX6 mis6-GFP:KanMX6 bqt1::hygMX6 | JCF10128 | (Fennell et al., 2015) |

**Supplementary Table 2**. **Nuclear morphological analysis;** for each of the strains studied, the averages and standard deviations calculated for each time series associated with a single-cell experiment. Five descriptors, area, axis length (and associated aspect ratios), circularity and convexity are indicated in their respective units.

| Strain | n | Area (μm$^2$) | Major (μm) | Minor (μm) | Circularity | Aspect ratio | Convexity |
|---|---|---|---|---|---|---|---|
| *wt* | 14 | 5.20 ± 1.88 | 4.09 ± 2.79 | 3.11 ± 1.18 | 0.16 ± 0.05 | 0.72 ± 0.29 | 0.74 ± 0.05 |
| *dhc1Δ* | 10 | 5.24 ± 2.56 | 5.98 ± 3.08 | 2.90 ± 1.09 | 0.12 ± 0.05 | 0.48 ± 0.35 | 0.70 ± 0.05 |
| *hrs1Δ* | 12 | 7.04 ± 4.95 | 6.03 ± 3.33 | 3.48 ± 1.37 | 0.12 ± 0.04 | 0.58 ± 0.41 | 0.71 ± 0.05 |
| *bqt1Δ* | 9 | 5.35 ± 2.81 | 4.25 ± 1.87 | 2.25 ± 0.32 | 0.12 ± 0.04 | 0.52 ± 0.28 | 0.76 ± 0.03 |
| *rdh54Δ* | 9 | 9.89 ± 3.56 | 4.25 ± 0.99 | 2.89 ± 0.54 | 0.10 ± 0.03 | 0.68 ± 0.34 | 0.73 ± 0.03 |

**Supplementary Table 3**. **Nuclear motion segment duration and spectral richness;** for each of the strains studied, time in minutes that, in average and with its associated standard deviation (or as the lower bound) last each of the stages deriving from significant segmentation. The number of samples for each case is indicated at the single-cell level, always from at least three repetitions. Dominant periods different than infinity are indicated when possible; associated Power Spectral Density (PSD) is established as the maximum observed for a specific dataset, considering the entire time-series sequence. Whole dots (•) indicate non-availability of data.

| Strain | n | Segment duration (minutes) | | | Spectral richness | |
|---|---|---|---|---|---|---|
| (Chr) | | ***High*** | ***Medium*** | ***Low*** | $p_1$ **(minutes)** | **PSD (**$p_1$**)** |
| *wt* | 14 | > 110 | 33 ± 4 | 50 ± 12 | 6.25 ± 2.09 | 6425.87 |
| *dhc1Δ* | 10 | > 70 | 29 ± 7 | 88 ± 12 | ∞ | 9.21 |
| *hrs1Δ* | 12 | > 91 | 28 ± 6 | 71 ± 15 | 9.61 ± 4.19 | 6.31 |
| *bqt1Δ* | 9 | • | • | • | ∞ | 175.43 |
| *rdh54Δ* | 9 | > 51 | 34 ± 7 | 91 ± 9 | 5.71 ± 1.74 | 11106.28 |

**Supplementary Table 4**. **SPB trajectory analysis: segment duration and spectral richness;** for each of the strains studied, time in minutes that, in average and with its associated standard deviation (or as the lower bound) last each of the stages deriving from significant segmentation. The number of samples for each case is indicated at the single-cell level, always from at least three repetitions. Power Spectral Density (PSD) is the maximum observed for a specific dataset.

| **Strain** | **n** | **Segment duration (minutes)** | | | **Spectral richness** | |
|---|---|---|---|---|---|---|
| (SPB) | | ***High*** | ***Medium*** | ***Low*** | $\boldsymbol{p_1}$ **(minutes)** | **PSD(**$\boldsymbol{p_1}$**)** |
| *wt* | 9 | > 71 | 29 ± 8 | 31 ± 6 | 6.21 ± 2.62 | 3854.90 |
| *dhc1*∆ | 9 | > 81 | 34 ± 3 | 60 ± 11 | ∞ | 10319.49 |
| *hrs1*∆ | 11 | > 50 | 35 ± 3 | 48 ± 8 | 6.25 ± 1.57 | 3.82 |
| *bqt1*∆ | 12 | > 75 | 37 ± 4 | 47 ± 6 | 6.85 ± 4.18 | 33657.05 |
| *rdh54*∆ | 11 | > 91 | 36 ± 7 | 51 ± 13 | 5.53 ± 3.14 | 7551.96 |
